# *N*-glycolylneuraminic acid serum biomarker levels are elevated in breast cancer patients at all stages of disease

**DOI:** 10.1101/2021.06.21.449179

**Authors:** Lucy K. Shewell, Christopher J. Day, Jamie R. Kutasovic, Jodie L. Abrahams, Jing Wang, Jessica Poole, Colleen Niland, Kaltin Ferguson, Jodi M. Saunus, Sunil R. Lakhani, Mark von Itzstein, James C. Paton, Adrienne W. Paton, Michael P Jennings

**Affiliations:** Institute for Glycomics, Griffith University, Gold Coast, Australia; UQ Centre for Clinical Research, Faculty of Medicine, The University of Queensland, Herston, QLD Australia; Pathology Queensland, The Royal Brisbane and Women’s Hospital, Herston, QLD Australia; Research Centre for Infectious Diseases, Department of Molecular and Biomedical Science, University of Adelaide, Adelaide, Australia; Glycosciences Laboratory, Department of Metabolism, Digestion and Reproduction, Imperial College London, London, UK

**Keywords:** *N*-glycolylneuraminic acid, Neu5Gc, biomarker, breast cancer, diagnostic

## Abstract

**Background:** Normal human tissues do not express glycans terminating with the sialic acid *N*-glycolylneuraminic acid (Neu5Gc), yet Neu5Gc-containing glycans have been consistently found in human tumor tissues, cells and secretions and have been proposed as a cancer biomarker. We engineered a Neu5Gc-specific lectin called SubB2M, and previously reported elevated Neu5Gc biomarkers in serum from ovarian cancer patients using a Surface Plasmon Resonance (SPR)-based assay. Here we report an optimized SubB2M SPR-based assay and use this new assay to analyse sera from breast cancer patients for Neu5Gc levels.

**Methods:** To enhance specificity of our SPR-based assay, we included a non-sialic acid binding version of SubB, SubB_A12_, to control for any non-specific binding to SubB2M, which improved discrimination of cancer-free controls from early-stage ovarian cancer. We analysed 96 serum samples from breast cancer patients at all stages of disease compared to 22 cancer-free controls using our optimized SubB2M-_A12_-SPR assay. We also analysed a collection of serum samples collected at 6 monthly intervals from breast cancer patients at high risk for disease recurrence or spread.

**Results:** Analysis of sera from breast cancer cases revealed significantly elevated levels of Neu5Gc biomarkers at all stages of breast cancer. We show that Neu5Gc serum biomarker levels can discriminate breast cancer patients from cancer-free individuals with 98.96% sensitivity and 100% specificity. Analysis of serum collected prospectively, post-diagnosis, from breast cancer patients at high risk for disease recurrence showed a trend for a decrease in Neu5Gc levels immediately following treatment for those in remission.

**Conclusions:** Neu5Gc serum biomarkers are a promising new tool for early detection and disease monitoring for breast cancer that may complement current imaging- and biopsy-based approaches.

## Background

Aberrant glycosylation is one of the hallmarks of cancer cells. Normal human tissues do not express glycans terminating with the sialic acid *N*-glycolylneuraminic acid (Neu5Gc) as humans express an inactive cytidine monophosphate *N*-acetylneuraminic acid (Neu5Ac) hydroxylase (CMAH) enzyme (1, 2); the only enzyme known to convert Neu5Ac to Neu5Gc. Nevertheless, Neu5Gc-containing glycans have been consistently found in human tumor tissues, cells and secretions (3–10), and have been proposed as a tumor biomarker (6, 11, 12).

Little progress has been made towards the development of a Neu5Gc biomarker-based assay for cancer detection and patient monitoring due to the lack of sufficiently sensitive and specific tools to detect this potential glyco-marker in a clinically relevant biological sample. We have improved upon the current methods for the detection of Neu5Gc (13–15) by developing a lectin with enhanced sensitivity and specificity for this glycan in the context of complex biological samples. This new lectin is derived from the B-subunit of the Shiga toxigenic *Escherichia coli* (STEC) Subtilase cytotoxin (SubAB), which recognizes α2-3 linked Neu5Gc (16). We used structure aided design to engineer this lectin to ablate the recognition of Neu5Ac and to expand the recognition from only α2-3-linked Neu5Gc to include both α2-3 and α2-6 Neu5Gc linkages to substituent sugars (17). This improved Neu5Gc-specific lectin was called SubB2M (17, 18). In a SPR-based SubB2M assay, we previously reported that the serum of ovarian cancer patients at all stages of disease has elevated levels of Neu5Gc-containing biomarkers compared to cancer-free females (19). This demonstrated the potential utility of Neu5Gc-containing biomarkers in the early detection of ovarian cancer, for which there is currently no universally applicable blood-based biomarker. The best currently available biomarker for ovarian cancer is the human cancer antigen 125 (CA125), also known as MUC16, a heavily glycosylated mucin (20). Serum CA125 levels are elevated in approximately 80% of ovarian cancer cases at the time of diagnosis (21). However, CA125 serum levels may also be elevated in non-malignant conditions including pregnancy, endometriosis, ovarian cysts, pelvic inflammatory disease and in the follicular phase of the menstrual cycle (22). As a screening tool for ovarian cancer, longitudinal measurement of CA125 levels has been shown to improve sensitivity and specificity for early detection. However, outcomes from the largest ovarian cancer screening trial to date, the UK Collaborative Trial of Ovarian Cancer Screening (UKCTOCS) show that screening including CA125 did not significantly reduce mortality (23, 24). Hence CA125 is currently the only approved ovarian cancer serum biomarker, limited to monitoring response to therapy and disease recurrence in post-menopausal women (23).

Breast cancer is the most frequently diagnosed cancer among women worldwide and is the leading cause of cancer death in over 100 countries (25). Detection of breast cancer at the early stages is associated with better patient outcomes including lower morbidity and lower mortality rates (26). Mammography is currently the main screening tool for the early detection of breast cancer; however, this method has limitations. For example, the sensitivity of mammography in women with dense breasts is reduced from 85 % to 47-64 % (27), yet these women have an increased risk of developing breast cancer (28). Cancer antigen 15-3 (CA 15-3) is the most widely used serum biomarker for breast cancer, and is approved for monitoring treatment efficacy only, due to the low sensitivity in early detection (29, 30). CA 15-3 is a secreted form of MUC1, a heavily glycosylated mucin (31, 32). High levels of circulating CA 15-3 have been found in breast cancer patients (33); however, like CA125 for ovarian cancer, levels of serum CA 15-3 are also elevated in other physiological conditions, such as pregnancy (34) and coronary heart disease (35). Despite decades of research, there is no single serum biomarker that has proved useful for the early detection or monitoring of recurrence in breast cancer (36, 37).

In this study we developed an improved SubB2M-based SPR assay methodology and used this assay to analyze serum samples collected from breast cancer patients to determine whether detection of Neu5Gc biomarkers may be relevant in screening for and monitoring of breast cancer.

## Methods

### Expression and purification of SubB2M and SubB_A12_

The recombinant SubB2M and SubB_A12_ proteins were expressed and purified as previously described (17, 38). Briefly, SubB2M and SubB_A12_ were expressed in *E. coli* BL21 (DE3) cells transformed with the SubB2M or SubB_A12_ expression constructs, respectively, as His6-tagged fusion proteins, which were then purified by Ni-NTA affinity chromatography.

### Glycan array analysis of SubB2M and SubB_A12_

Neu5Ac/Neu5Gc glycan array slides were purchased from Z-Biotech (Aurora, Colorado, USA). A 16-subarray slide array was used and glycan array analysis of SubB2M and SubB_A12_ was performed as described previously (18) and as described in Supplementary Table 1. The full list of glycans on the array can be found at https://www.zbiotech.com/neu5gc-neu5ac-n-glycan-array.html.

### Development and use of the SubB2M-_A12_-SPR assay for Neu5Gc serum biomarkers

SPR was conducted using the Biacore S200 system (GE) with immobilization of SubB2M and SubB_A12_ performed essentially as described previously (19). For glycan analysis, SubB2M was immobilized through flow cells 2 and 3 and SubB_A12_ was immobilized through flow cell 4 (capture levels: 5000-6000 Response Units (RU) onto a series S sensor chip CM5 (GE) using the NHS capture kit. Glycans purchased from Chemily Glycoscience (Atlanta, GA) were analysed across a five-fold dilution series in PBS at a maximum concentration of 20μM. Analysis was run using single cycle analysis and double reference subtraction on the Biacore S200 evaluation software. For glycoprotein analysis, SubB2M was immobilized through flow cells 2 and 3 and SubB_A12_ was immobilized through flow cell 4. Flow cell 1 was run as a blank immobilization. After immobilization, a start up cycle of 0.5 % normal human serum (Sigma-Aldrich, Cat No. H4522) was run over the immobilized SubB proteins for 10 steps of 30 seconds at 30 μL/minute flow rate to condition the chip. A final wash of 10 mM Tris/1 mM EDTA was run for 30 seconds at a 30 μL/minute flow rate prior to beginning the data collection. SPR analysis was performed using multi-cycle analysis and double reference (values from flow cell 1 and 0.5 % normal human serum only) subtraction using the Biacore S200 evaluation software. At least two independent SPR runs were performed for each sample set.

For analysis of human sera, samples were diluted 1:200 in PBS and analyzed in duplicate in each SPR run as described above. RU values obtained for each serum sample with SubB_A12_ (flow cell 4) were subtracted from the RU values obtained with SubB2M from flow cells 2 and 3 and averaged to obtain the final RUs used for conversion to GPUs. Two independent SPR runs were performed for each sample set.

### Development of Glycoprotein Units (GPUs) standard curve for normalization of data from the SubB2M-_A12_-SPR assays

SPR was conducted as described above with SubB2M immobilized through flow cells 2 and 3 and SubB_A12_ immobilized through flow cell 4. To generate an internal calibration curve, bAGP and CA125 were combined at starting concentrations of 15 μg/ml and 15 units/ml, respectively, in 0.5 % normal human serum. For further detail, see Supplementary Methods.

### Mass spectrometry glycomic analysis of standard glycoproteins

To confirm the presence of Neu5Gc on the glycoprotein standards (bAGP and human CA125) *N* and *O*-glycans were released and analysed by PGC-LC-MS/MS as previously described (39, 40). For further details, see Supplementary Methods.

### Human serum samples

#### Victorian Cancer Biobank samples

Serum samples from cancer-free (normal) females and serum samples from patients with Stage I (n=12), Stage II (n=11), Stage IIIC (n=10) and Stage IV ovarian cancer (n=14) were obtained from the Victorian Cancer Biobank and have been described previously (19). 24 serum samples from patients with Stage I breast cancer, 24 with Stage II breast cancer, 24 with Stage III breast cancer and 24 with Stage IV breast cancer were also obtained from the Victorian Cancer Biobank under application 17020. As described in our previous study (19), ‘normal’ controls are defined as patients with an apparent non-malignancy diagnosis at the time the sample is taken. The serum samples were collected immediately pre-operatively using Serum Separation Tubes (BD) and were processed and stored at −80 °C within two hours of collection. The patient data and serum samples used in this project were provided by the Victorian Cancer Biobank with informed consent from all donors and use of the samples was approved by the Griffith University HREC (GU Ref No: 2017/732) in accordance with the National Statement on Ethical Conduct in Human Research. The majority of the breast cancers in the cohort were the most common form of breast cancer, invasive ductal carcinoma. The remainder included 8 cases of invasive lobular carcinoma and 6 cases of mucinous carcinoma. Information for each of the ovarian cancer serum samples used in this study can be found in our previous publication (19) while information regarding each of the breast cancer serum samples can be found in Supplementary Table 2.

#### Circulating Biomarkers of Relapse in Breast Cancer (Circ.BR) cohort

Circ.BR was established in 2013 as part of the Brisbane Breast Bank (41). Patients with breast cancer who are at high risk for disease recurrence or spread (inclusion criteria below) are followed prospectively for 5 years, with serial collection of blood samples taken at 6 monthly intervals and tumor tissue collected at the time of surgery. Human research ethics committees of The University of Queensland (ref. 2005000785) and The Royal Brisbane and Women’s Hospital (2005/022) approved the study with written informed consent obtained from each subject. Serial blood samples from 9 patients who experienced a relapse (median 4 samples per patient, range 3-9) and 6 patients who were free from recurrence (median 7 samples per patient, range 6-8) at the time of the study were analyzed, with a median follow-up of 19.2 months for the relapse group, and 43.9 months for the recurrence-free group. Detailed information for each patient in the Circ.BR cohort are shown in Supplementary Table 3.

### Circ.BR inclusion criteria

Invasive breast cancer, grade 3 or grade 2 (score >6) invasive cancer and axillary lymph node positive; OR grade 1 and grade 2 (score ≤6) invasive cancer with adverse features such as tumor size >5cm; Family history (NBOCC group 3)/gene carrier; or previous history of breast cancer.

#### Statistical analysis

All statistical analyses were performed using GraphPad Prism 8.0. The mean GPUs between normal serum samples compared to cancer patient serum samples were analyzed by two-tailed, unpaired *t*-tests, with a *P* value of <0.05 considered significant. Optimal cut-off values from Receiver operating characteristics (ROC) analyses were determined by maximizing the sum of specificity and sensitivity.

## Results

### Development of the optimized SubB2M-_A12_-SPR assay for detection of Neu5Gc serum biomarkers

In a previous study, we analyzed serum samples from ovarian cancer patients with our SubB2M lectin via the highly sensitive method of SPR (19). This assay is based on label-free detection of Neu5Gc biomarkers that bind to the SubB2M lectin and elicit a response in SPR (see Figure 1). In this study we aimed to improve this SPR-based assay by including a parallel analysis of all samples with a non-sialic acid binding version of SubB called SubB_A12_. SubB_A12_ has the most critical amino acid residue of the B subunit required for binding Neu5Gc mutated from a serine to an alanine (Ser12 > Ala12) (16). Mutation of this Ser residue abolishes interactions with the C1 carboxylate group of sialic acid and thus the SubB_A12_ mutant cannot bind any sialylated glycans. Any binding to SubB_A12_ observed with serum samples must be due to non-sialic acid-dependent interactions of serum components with the SubB protein, for example the binding of antibodies that may recognize the SubB portion of the SubAB toxin. The lack of binding to sialylated glycans by the SubB_A12_ mutant has been described previously (16) and was further confirmed herein with an analysis of SubB_A12_ specificity using a Neu5Ac/Gc glycan microarray, where negligible binding was observed to either Neu5Ac or Neu5Gc-containing glycans by this mutant protein compared to SubB2M **(Table S1 and Figure S1)**. The kinetics of the interaction of SubB2M and SubB_A12_ using SPR analysis with a range of paired synthetic oligosaccharides presenting either a Neu5Ac or Neu5Gc are shown in **Figure 1A** and confirms the loss of all sialic acid binding by SubB_A12_. In the optimized SubB2M-_A12_-SPR assay for the analysis of serum samples, the SPR Response Units (RUs) detected with SubB_A12_ for each serum sample are subtracted from the RUs detected with SubB2M to control for any non-Neu5Gc-dependent binding of serum components to SubB2M **(Figure 1B)**.

**Figure 1.**
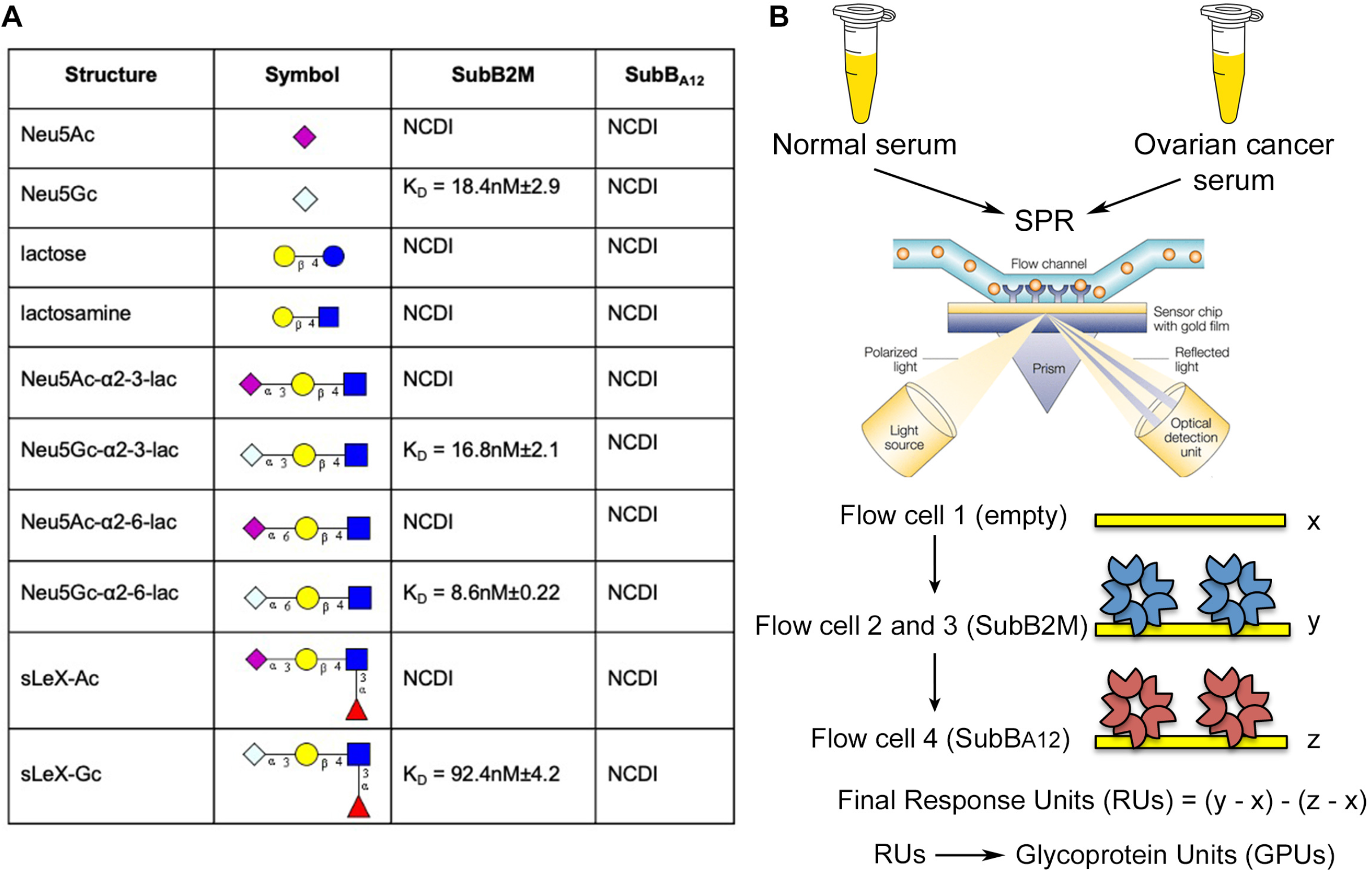
Characterization of the SubB2M-_A12_-SPR assay workflow used in this study. **A)** SPR analysis of Neu5Ac/Neu5Gc glycan pairs. NCDI: No concentration dependent interaction with glycan up to 20 μM. **B)** Optimized SubB2M-_A12_-SPR assay. Serum from cancer-free (normal) females and ovarian cancer patients were analyzed by SPR with SubB2M immobilized onto the surface of the sensor chip through flow cells 2 and 3 and SubB_A12_ immobilized onto the sensor surface through flow cell 4.

To further refine our SubB2M-_A12_-SPR assay we established a standard curve using a mixture of two commercially available Neu5Gc-containing glycoproteins at known concentrations to be included as an internal control in each SPR analysis to normalize all data from all studies to common units. As the identity and nature of the Neu5Gc glycoconjugates detected in the ovarian cancer patient serum samples from our 2018 study (19) is currently unknown, we selected a combination of glycoproteins representing a high molecular weight glycoprotein with low Neu5Gc glycosylation and a lower molecular weight glycoprotein with high Neu5Gc glycosylation. These glycoproteins were the human tumor antigen CA125 and bovine AGP (bAGP), respectively. We have previously confirmed the presence of Neu5Gc on bAGP (17) and CA125 (18), and this was reconfirmed for both of the control glycoproteins by mass spectrometry analysis **(Figure S2)**. The standard curve generated by this mixture of glycoproteins diluted into 0.5 % normal human serum was used to calibrate the SubB2M-_A12_-SPR assay. The response units (RUs) obtained for each serum sample was converted to Glycoprotein Units (GPUs) (representative standard curve shown in **Figure S3**).

In our 2018 ovarian cancer study (19) we analyzed serum samples from subjects in Stage I (n=12), Stage II (n=11), Stage IIIC (n=10) and Stage IV ovarian cancer (n=14) as well as serum samples from 22 cancer-free females. In the current study, we reanalyzed this sample set using our optimized SubB2M-_A12_-SPR assay **(Figure 2)**. Figure 2A shows the data before subtraction of non-specific binding to SubB_A12_ from each serum sample while Figure 2B shows the data after subtraction of SubB_A12_ responses. As we saw with our original analysis of this sample set (19), significantly elevated serum Neu5Gc biomarker levels were detected at all stages of ovarian cancer compared to cancer-free female controls.

**Figure 2.**
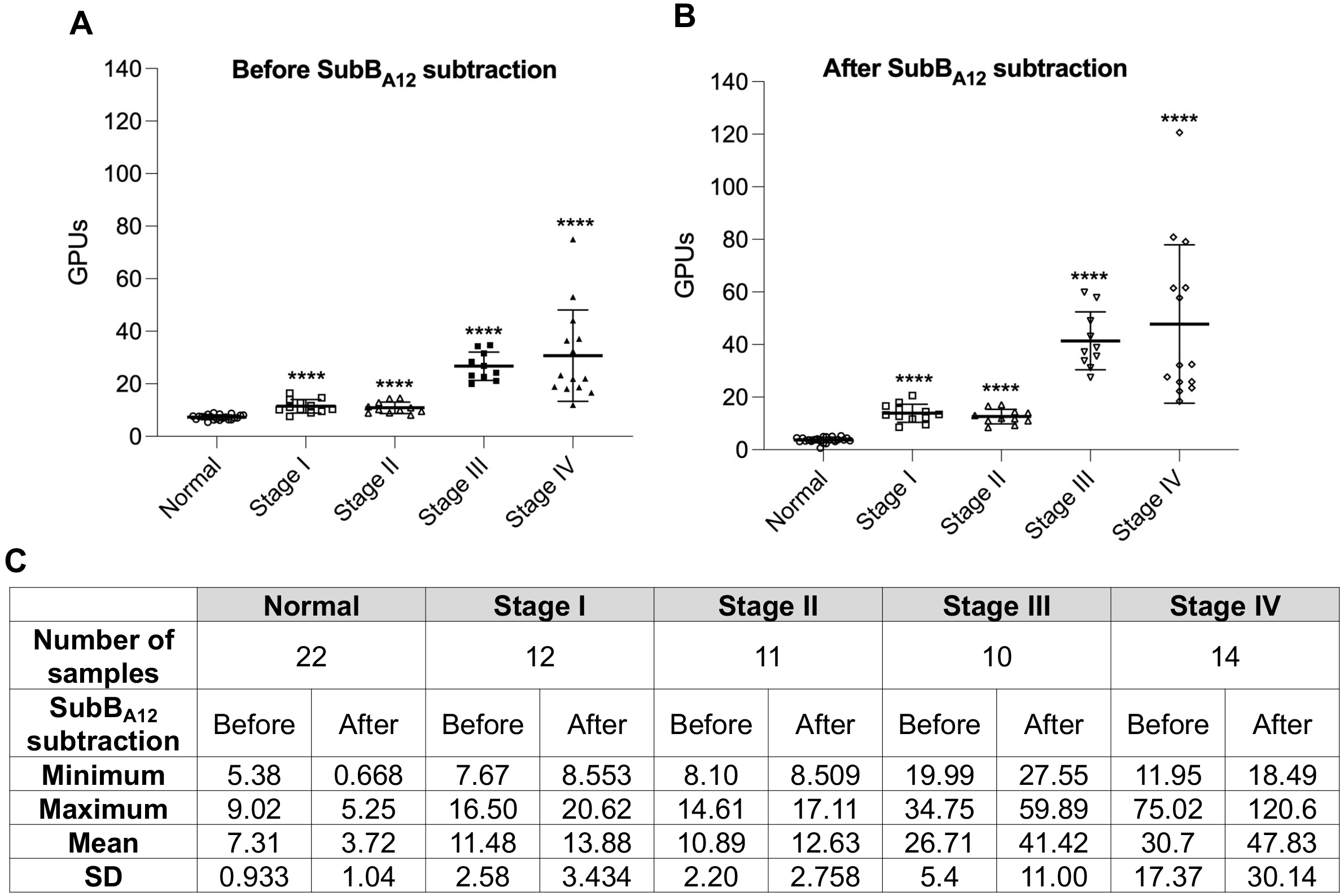
Analysis of cancer-free and Stage I-IV ovarian cancer patient serum samples with the optimized SubB2M-_A12_-SPR assay. 22 serum samples from cancer-free (normal) females, 12 patients with Stage I ovarian cancer, 11 with Stage II ovarian cancer, 10 with Stage IIIC ovarian cancer and 14 with Stage IV ovarian cancer were analyzed by the optimized SubB2M-SPR assay. The mean GPUs from duplicate analyses for each serum sample determined **A)** before and **B)** after subtraction of binding due to SubB_A12_ are shown. Error bars = ± 1 SD from the mean for each group. Statistical analysis was performed using two-tailed unpaired *t*-tests. **** = *P*-value <0.0001 compared to Normal. Duplicate, independent assays were performed with both showing the same trends. One representative assay is shown. **C)** Descriptive statistics of data from ovarian cancer patient serum samples and cancer-free controls.

Receiver-Operating Characteristic (ROC) analyses were performed on the serum Neu5Gc levels detected with the optimized SubB2M-_A12_-SPR assay before and after SubB_A12_ subtraction **(Supplementary Table 4, Figure S4)**. Subtraction of binding due to SubB_A12_ improved the ability of our SPR-based assay to distinguish cancer-free individuals from ovarian cancer patients at all stages to 100 % specificity and 100 % sensitivity.

### Serum Neu5Gc levels can discriminate breast cancer patients from cancer-free individuals with high specificity and sensitivity

We then used the refined SubB2M-_A12_-SPR assay to determine whether elevated levels of Neu5Gc biomarkers could be detected in serum from patients with breast cancer compared to cancer-free controls. We analyzed a collection of breast cancer serum samples across all stages of disease (24 Stage I, 24 Stage II, 24 Stage III and 24 Stage IV) with the same set of cancer-free females used for the ovarian cancer analyses. The SubB2M-_A12_-SPR analysis **(Figure 3)** shows that significantly elevated levels of Neu5Gc biomarkers were detected in serum samples from all stages of disease. Detailed clinical information for each of the serum samples can be found in Supplementary Table 2.

**Figure 3.**
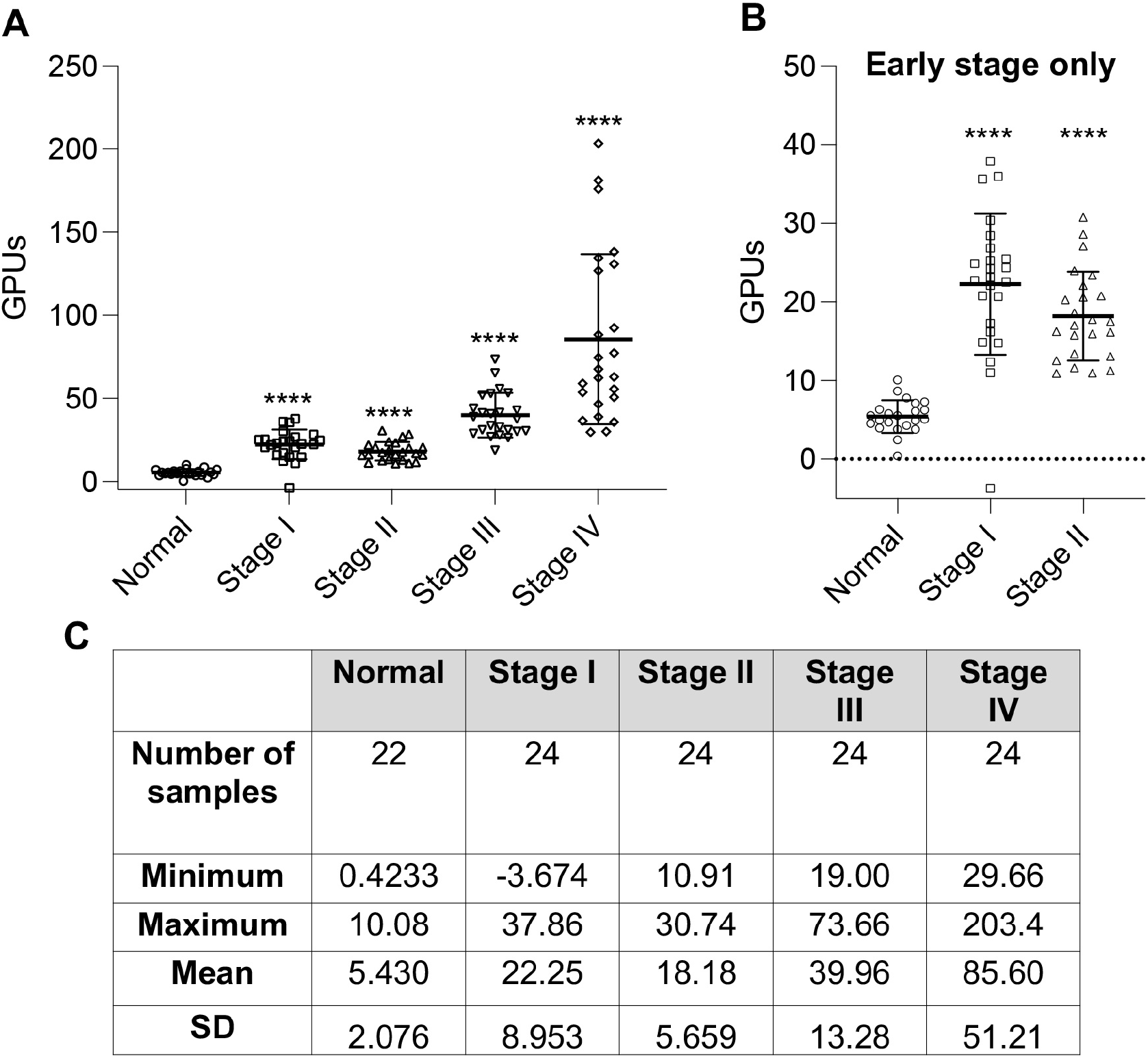
SubB2M-_A12_-SPR analysis of cancer-free and Stage I – IV breast cancer serum samples. **A)** 22 serum samples from cancer-free females, 24 Stage I, 24 Stage II, 24 Stage III and 24 Stage IV serum samples were analyzed by SubB2M-_A12_-SPR assay. The mean GPUs from duplicate analyses for each serum sample are shown. Error bars = ± 1 SD from the mean for each group. Statistical analysis was performed using two-tailed unpaired *t*-tests. **** = *P*-value <0.0001 compared to Normal. Duplicate, independent assays were performed with both showing the same trends. One representative assay is shown. Only values above 0 are shown. **B)** Cancer-free controls compared to early stage breast cancer samples only (Stage I and II samples only from Figure 3A). **C)** Descriptive statistics of data from breast cancer patient serum samples and cancer-free controls.

ROC analyses were performed to assess the ability of serum Neu5Gc levels detected with the optimized SubB2M-_A12_-SPR assay to discriminate cancer-free females from patients from each stage of breast cancer (**Supplementary Table 5, Figure S5**). When all stages are considered as one group with an optimal ROC cut-off value (>10.49 GPUs), as would be the case for a diagnostic screen, the SubB2M-_A12_-SPR assay has 98.96% sensitivity and 100% specificity to distinguish patients with breast cancer across all stages of disease from cancer-free individuals (**Figure 4**). In summary, this test achieves 100% specificity and 100% sensitivity for patients with Stage II-IV disease, however, due to one individual data point in the Stage I group, below the limit of detection of our assay, the overall sensitivity did not reach 100%. These data indicate that the detection of serum Neu5Gc-biomarkers with SubB2M has potential to detect breast cancer at all stages of disease.

**Figure 4.**
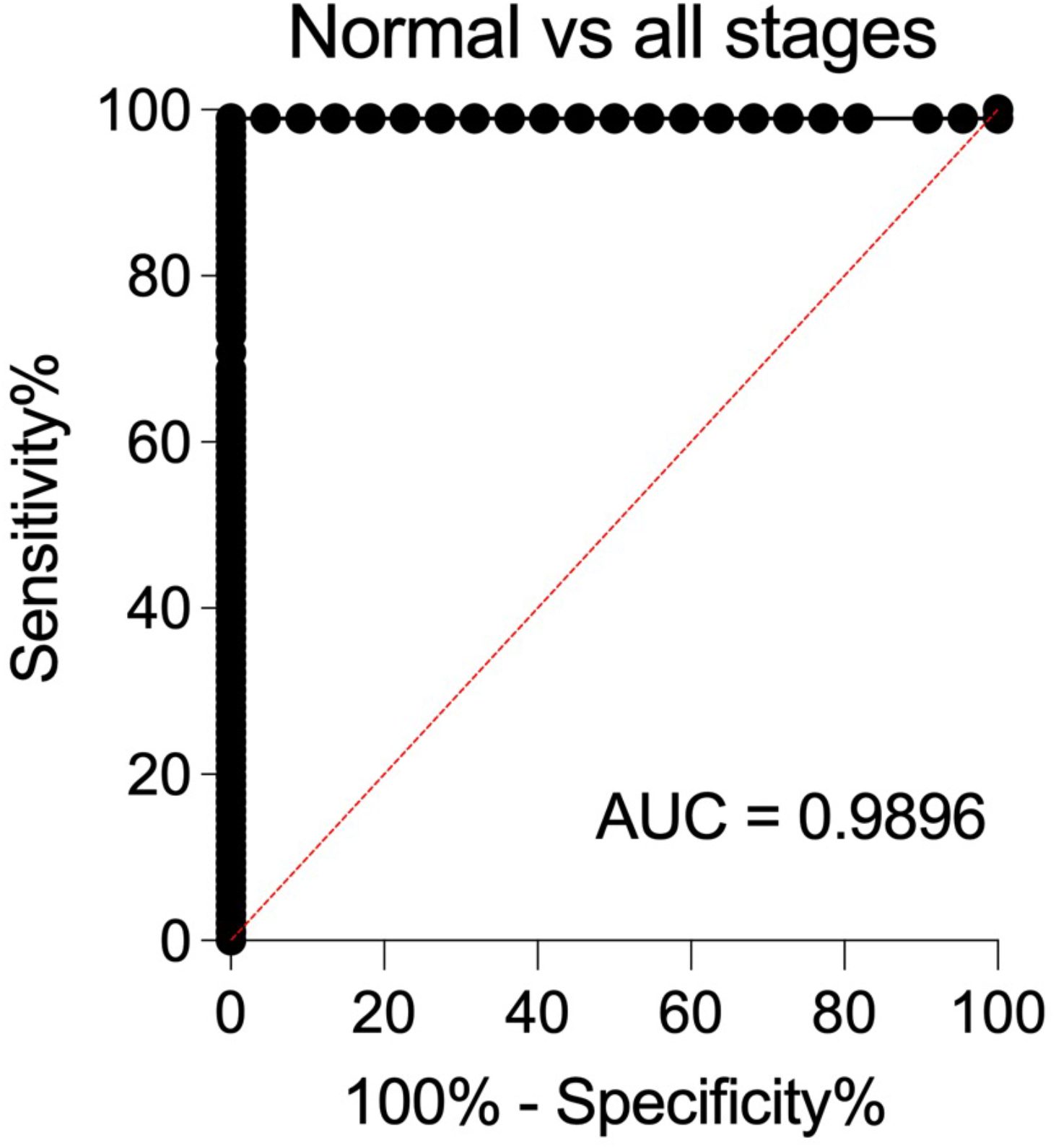
ROC curve depicting the ability of serum Neu5Gc levels determined by the optimized SubB2M-_A12_-SPR assay to distinguish breast cancer patients from cancer-free (normal) individuals. Sensitivity% (true positive rate; ability to detect disease) is plotted against 100 %-specificity% (false positive rate or 100 %-true negative rate; ability to detect lack of disease). ROC analyses were performed with the data shown in Figure 3A using Graphpad Prism 8.0.

### Analysis of serum Neu5Gc levels using SubB2M has potential utility for treatment monitoring in breast cancer

The Circ.BR cohort is a collection of breast cancer patients with serum samples collected at 6 monthly intervals, allowing us the opportunity to analyze Neu5Gc biomarker levels over the course of disease in these patients at high risk for disease recurrence or spread. Detailed clinical information for each patient can be found in Supplementary Table 3. Analysis of the available serum samples from 15 cases (6 cases in remission, 9 cases with relapse) from this cohort showed a trend for a decrease in Neu5Gc levels immediately following the first line of treatment in the cases who did not have a tumor recurrence, but only in some of the recurrence cases. Figure 5 shows a representative plot from one remission case and one relapse case, with the remaining cases shown in Supplementary Figure S6. These data demonstrate the potential utility of the assessment of serum Neu5Gc levels to monitor treatment response during breast cancer and warrants further exploration and validation.

**Figure 5.**
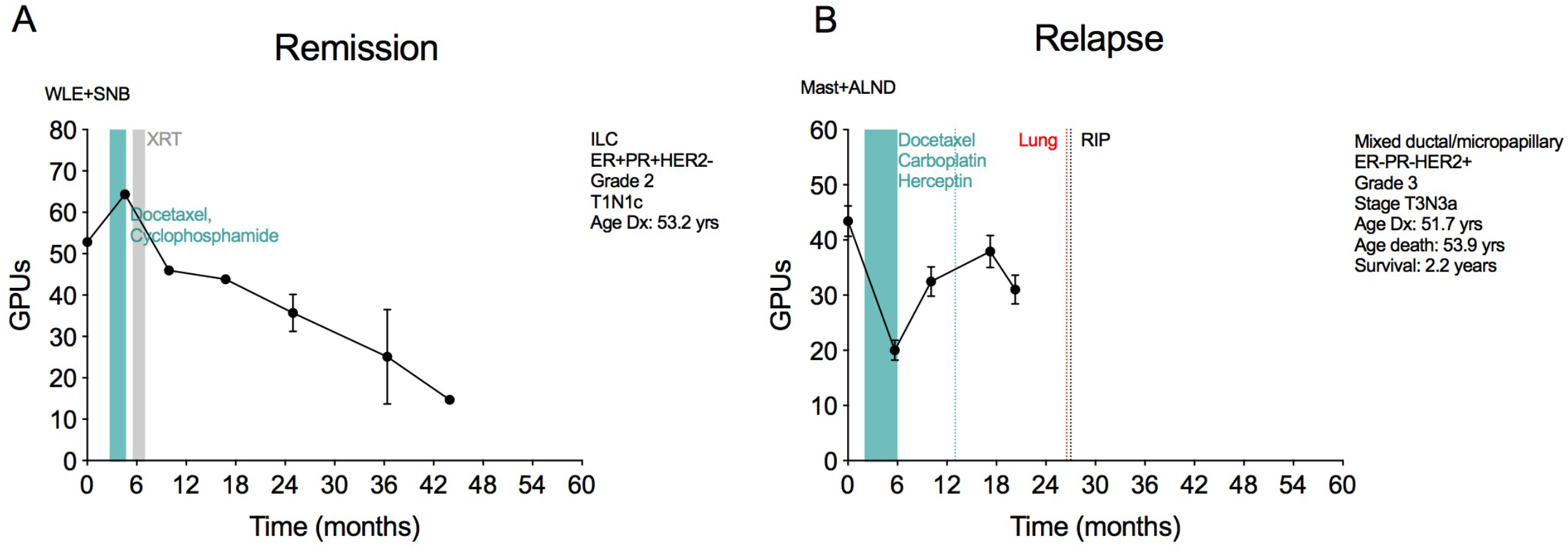
Representative plots of serum Neu5Gc levels determined by SubB2M-_A12_-SPR assay for A) one remission case and B) one relapse case from the Circ.BR cohort. The mean GPUs from duplicate analyses for each serum sample are shown. Error bars = ± 1 SD from the mean for each group. Two independent assays were performed with both showing the same trends with results from one assay presented. Plots for the remaining cases are shown in Supplementary Figure S6. Clinical information for each patient is shown in the top right of each plot with treatment history and metastases overlaid. ALND: Axillary lymph node dissection, ILC: Invasive Lobular Carcinoma, Mast: mastectomy, SNB: sentinel node biopsy, WLE: wide local excision, XRT: radiation therapy.

## Discussion

CA125 is currently the best performing serum biomarker for ovarian cancer, but due to its limitations, it is not currently used as a screening tool (23). Using our new, optimized SubB2M-_A12_-SPR assay, we are able to distinguish cancer-free females from ovarian cancer patients at all stages of disease with 100 % specificity and 100 % specificity.

The SubB2M-_A12_-SPR assay is an improved version of our previously described SubB2M-SPR assay (19). Firstly, we included subtraction of non-specific binding of serum components to SubB_A12_. The inclusion of the SubB_A12_ mutant improved the specificity and sensitivity of our assay to distinguish cancer-free females from early-stage ovarian cancer patients. Secondly, we now report Neu5Gc biomarker levels as GPUs. Including a standard curve in each SPR analysis allowed us to convert Neu5Gc biomarker levels in patient serum samples to GPUs to control for assay-to-assay variability and to accurately compare data from independent studies.

While screening mammography is used worldwide for early detection of breast cancer, this method has limitations, including accessibility for women in low-income countries (42), and there is currently no single serum biomarker for the early detection of breast cancer. Our analysis of 96 breast cancer patient serum samples from Stages I-IV, showing that serum Neu5Gc levels can distinguish patients with early-stage breast cancer from cancer-free individuals with high specificity and high sensitivity, and has perfect discriminative ability to distinguish later stage breast cancer from cancer-free individuals, indicates that SubB2M detection of serum Neu5Gc levels has the potential to be developed into a tool for the early detection of breast cancer. It is important to note that this cohort, whilst comprising mostly invasive ductal carcinoma, also included the less common invasive lobular carcinoma, and six cases of mucinous carcinoma. Mucinous carcinoma of the breast is a rare form of invasive ductal carcinoma, making up 2-3% of breast cancer patients. That all of these cancers could be differentiated from cancer-free controls by our assay supports the broad applicability of the approach. Our analysis of serially collected serum samples from the Circ.BR cohort showed that SubB2M detection of Neu5Gc levels also has potential to be used to monitor response to treatment, although further investigation of its utility, using a larger cohort, is required.

Neu5Gc is well known to be found in human cancer (in cells, tissues and secretions) (3, 5–9, 43), and may be found in trace amounts in some healthy human tissues (44). The source of Neu5Gc in humans is assumed to be dietary incorporation following consumption of animal-derived food. Dietary acquisition of Neu5Gc and metabolic incorporation into human cells forms the basis of the ‘xenosialitis’ hypothesis, which proposes that autoantibodies against Neu5Gc at the cell surface promote inflammation and pathology (see reference 45 for a recent review) (45). The direct biochemical evidence for dietary uptake and expression in humans involved three human subjects consuming Neu5Gc in the form of porcine submaxillary mucin (44). Alternatively, It has been proposed that human cancer cells can produce endogenous Neu5Gc by providing electron donors that circumvent the CMAH inactivating truncation found in the human enzyme (46). The current study and our previous data (19) report elevated levels of Neu5Gc in serum can differentiate breast and ovarian cancer patients (n=143) from cancer-free controls, and supports the hypothesis of endogenous production of this biomarker by tumor cells in addition to potential dietary uptake.

Our development of the purpose-engineered SubB2M lectin implemented in the SubB2M-_A12_-SPR assay provides a sensitive, specific and practical reagent for the measurement of Neu5Gc-containing glycoproteins in serum, which are an under-explored class of circulating tumor biomarkers. The detection of Neu5Gc glycosylation via SubB2M may also be used in conjunction with existing biomarker assays to improve specificity and sensitivity and represents a promising area for future studies.

## Conclusions

Minimally invasive, rapid and reliable methods for the early detection of ovarian and breast cancer will have a significant impact on patient survival rates. Biomarkers that can accurately and specifically monitor response to treatment and disease progression would also drastically improve patient outcomes. Our SubB2M lectin assay offers the opportunity to exploit measurement of Neu5Gc biomarkers as a path to achieve these outcomes.

## Declarations

### Ethics approval and consent to participate

The patient data and serum samples used in this project were provided by the Victorian Cancer Biobank with informed consent from all donors and use of the samples was approved by the Griffith University HREC (GU Ref No: 2017/732). Human research ethics committees of The University of Queensland (ref. 2005000785) and The Royal Brisbane and Women’s Hospital (2005/022) approved the Circ.BR study with written informed consent obtained from each subject.

### Consent for publication

Not applicable

### Availability of data and materials

All data generated or analysed during this study are included in this published article and its supplementary information files.

### Author’s contributions

LKS curated all serum samples, analysed and interpreted SPR data, produced figures, contributed to design of the study and was a major contributor to writing of the manuscript, CJD acquired and analysed SPR and glycan array data, produced figures, contributed to conceptualisation and design of the study and contributed to writing of the manuscript, JRK contributed to analysing and interpreting Circ.BR data and writing of the manuscript, JLA acquired and interpreted mass spectrometry data and contributed to review and editing of the manuscript, JW contributed to curation of serum samples, SPR analysis and review and editing of the manuscript, JP acquired glycan array data and contributed to review and editing of the manuscript, CN contributed to the Circ.BR cohort and read and approved the final manuscript, KF contributed to the Circ.BR cohort and read and approved the final manuscript, JMS contributed to the Circ.BR cohort and read and approved the final manuscript SRL contributed to the Circ.BR cohort and read and approved the final manuscript, MvI provided resources and read and approved the final manuscript, JCP contributed to conceptualisation and design of the study, provided SubB2M and SubB_A12_ proteins and reviewed and edited the manuscript, AWP contributed to conceptualisation and design of the study, provided SubB2M and SubB_A12_ proteins and reviewed and edited the manuscript, MPJ conceptualised and designed the study, contributed to writing of the manuscript and provided supervision of the study.

## Acknowledgements

We thank the Victorian Cancer Biobank for providing all patient data and serum samples. The Victorian Cancer Biobank, through the Cancer Council Victoria as Lead Agency, is supported by the Victorian Government through the Victorian Cancer Agency, a business unit of the Department of Health and Human Services.

## References

1 Varki NM, Varki A. Diversity in cell surface sialic acid presentations: implications for biology and disease. Lab Invest. 2007;87(9):851–857.

2 Chou HH, Hayakawa T, Diaz S, et al. Inactivation of CMP-N-acetylneuraminic acid hydroxylase occurred prior to brain expansion during human evolution. Proceedings of the National Academy of Sciences of the United States of America. 2002;99(18):11736–11741.

3 Inoue S, Sato C, Kitajima K. Extensive enrichment of N-glycolylneuraminic acid in extracellular sialoglycoproteins abundantly synthesized and secreted by human cancer cells. Glycobiology. 2010;20(6):752–762.

4 Samraj AN, Laubli H, Varki N, Varki A. Involvement of a non-human sialic Acid in human cancer. Front Oncol. 2014;4:33.

5 Higashi H, Hirabayashi Y, Fukui Y, et al. Characterization of N-glycolylneuraminic acid-containing gangliosides as tumor-associated Hanganutziu-Deicher antigen in human colon cancer. Cancer research. 1985;45(8):3796–3802.

6 Malykh YN, Schauer R, Shaw L. N-Glycolylneuraminic acid in human tumours. Biochimie. 2001;83(7):623–634.

7 Furukawa J, Tsuda M, Okada K, et al. Comprehensive Glycomics of a Multistep Human Brain Tumor Model Reveals Specific Glycosylation Patterns Related to Malignancy. PloS one. 2015;10(7):e0128300.

8 Tzanakakis GN, Nikitovic D, Katonis P, Kanakis I, Karamanos NK. Expression and distribution of N-acetyl and N-glycolylneuraminic acids in secreted and cell-associated glycoconjugates by two human osteosarcoma cell lines. Biomedical chromatography : BMC. 2007;21(4):406–409.

9 Tzanakakis GN, Syrokou A, Kanakis I, Karamanos NK. Determination and distribution of N-acetyl- and N-glycolylneuraminic acids in culture media and cell-associated glycoconjugates from human malignant mesothelioma and adenocarcinoma cells. Biomedical chromatography : BMC. 2006;20(5):434–439.

10 Labrada M, Dorvignit D, Hevia G, et al. GM3(Neu5Gc) ganglioside: an evolution fixed neoantigen for cancer immunotherapy. Seminars in Oncology. 2018;45(1):41–51.

11 Marquina G, Waki H, Fernandez LE, et al. Gangliosides expressed in human breast cancer. Cancer research. 1996;56(22):5165–5171.

12 Higashi H, Sasabe T, Fukui Y, Maru M, Kato S. Detection of gangliosides as N-glycolylneuraminic acid-specific tumor-associated Hanganutziu-Deicher antigen in human retinoblastoma cells. Japanese journal of cancer research : Gann. 1988;79(8):952–956.

13 Priego-Capote F, Orozco-Solano MI, Calderon-Santiago M, Luque de Castro MD. Quantitative determination and confirmatory analysis of N-acetylneuraminic and N-glycolylneuraminic acids in serum and urine by solid-phase extraction on-line coupled to liquid chromatography-tandem mass spectrometry. Journal of chromatography A. 2014;1346:88–96.

14 Kitajima K, Varki N, Sato C. Advanced Technologies in Sialic Acid and Sialoglycoconjugate Analysis. Topics in current chemistry. 2015;367:75–103.

15 Diaz SL, Padler-Karavani V, Ghaderi D, et al. Sensitive and specific detection of the non-human sialic Acid N-glycolylneuraminic acid in human tissues and biotherapeutic products. PloS one. 2009;4(1):e4241.

16 Byres E, Paton AW, Paton JC, et al. Incorporation of a non-human glycan mediates human susceptibility to a bacterial toxin. Nature. 2008;456(7222):648–652.

17 Day CJ, Paton AW, Higgins MA, et al. Structure aided design of a Neu5Gc specific lectin. Scientific reports. 2017;7(1):1495.

18 Wang J, Shewell LK, Paton AW, Paton JC, Day CJ, Jennings MP. Specificity and utility of SubB2M, a new N-glycolylneuraminic acid lectin. Biochemical and biophysical research communications. 2018;500(3):765–771.

19 Shewell LK, Wang JJ, Paton JC, Paton AW, Day CJ, Jennings MP. Detection of N-glycolylneuraminic acid biomarkers in sera from patients with ovarian cancer using an engineered N-glycolylneuraminic acid-specific lectin SubB2M. Biochemical and biophysical research communications. 2018;507(1–4):173–177.

20 Yin BW, Lloyd KO. Molecular cloning of the CA125 ovarian cancer antigen: identification as a new mucin, MUC16. The Journal of biological chemistry. 2001;276(29):27371–27375.

21 Bast RC, Jr., Klug TL, St John E, et al. A radioimmunoassay using a monoclonal antibody to monitor the course of epithelial ovarian cancer. The New England journal of medicine. 1983;309(15):883–887.

22 Goonewardene T, Hall MR, Rustin GJS. Management of asymptomatic patients on follow-up for ovarian cancer with rising CA-125 concentrations. Lancet Oncology. 2007;8:813–821.

23 Jacobs I, Menon U, Ryan Aea. Ovarian cancer screening and mortality in the UK Collaborative Trial of Ovarian Cancer Screening (UKCTOCS): a randomised controlled trial. . Lancet. 2016;387:945–956.

24 Menon U, Gentry-Maharaj A, Burnell M, et al. Ovarian cancer population screening and mortality after long-term follow-up in the UK Collaborative Trial of Ovarian Cancer Screening (UKCTOCS): a randomised controlled trial. Lancet. 2021.

25 Bray F, Ferlay J, Soerjomataram I, Siegel RL, Torre LA, Jemal A. Global cancer statistics 2018: GLOBOCAN estimates of incidence and mortality worldwide for 36 cancers in 185 countries. CA: a cancer journal for clinicians. 2018;68(6):394–424.

26 Groot MT, Baltussen R, Uyl-de Groot CA, Anderson BO, Hortobagyi GN. Costs and health effects of breast cancer interventions in epidemiologically different regions of Africa, North America, and Asia. The breast journal. 2006;12 Suppl 1:S81–90.

27 Kolb TM, Lichy J, Newhouse JH. Comparison of the performance of screening mammography, physical examination, and breast US and evaluation of factors that influence them: an analysis of 27,825 patient evaluations. Radiology. 2002;225(1):165–175.

28 Boyd NF, Guo H, Martin LJ, et al. Mammographic density and the risk and detection of breast cancer. The New England journal of medicine. 2007;356(3):227–236.

29 Cheung KL, Graves CR, Robertson JF. Tumour marker measurements in the diagnosis and monitoring of breast cancer. Cancer treatment reviews. 2000;26(2):91–102.

30 Harris L, Fritsche H, Mennel R, et al. American Society of Clinical Oncology 2007 update of recommendations for the use of tumor markers in breast cancer. Journal of clinical oncology : official journal of the American Society of Clinical Oncology. 2007;25(33):5287–5312.

31 Gendler SJ, Lancaster CA, Taylor-Papadimitriou J, et al. Molecular cloning and expression of human tumor-associated polymorphic epithelial mucin. The Journal of biological chemistry. 1990;265(25):15286–15293.

32 Nath S, Mukherjee P. MUC1: a multifaceted oncoprotein with a key role in cancer progression. Trends in molecular medicine. 2014;20(6):332–342.

33 Devine PL, McGuckin MA, Ramm LE, Ward BG, Pee D, Long S. Serum mucin antigens CASA and MSA in tumors of the breast, ovary, lung, pancreas, bladder, colon, and prostate. A blind trial with 420 patients. Cancer. 1993;72(6):2007–2015.

34 Han SN, Lotgerink A, Gziri MM, Van Calsteren K, Hanssens M, Amant F. Physiologic variations of serum tumor markers in gynecological malignancies during pregnancy: a systematic review. BMC medicine. 2012;10:86.

35 Li X, Xu Y, Zhang L. Serum CA153 as biomarker for cancer and noncancer diseases. Progress in molecular biology and translational science. 2019;162:265–276.

36 Tang SS, Gui GP. Biomarkers in the diagnosis of primary and recurrent breast cancer. Biomarkers in medicine. 2012;6(5):567–585.

37 Loke SY, Lee ASG. The future of blood-based biomarkers for the early detection of breast cancer. European journal of cancer. 2018;92:54–68.

38 Paton AW, Srimanote P, Talbot UM, Wang H, Paton JC. A new family of potent AB(5) cytotoxins produced by Shiga toxigenic Escherichia coli. The Journal of experimental medicine. 2004;200(1):35–46.

39 Jensen P, Karlsson N, Kolarich D, Packer N. Structural analysis of N- and O-glycans released from glycoproteins. Nat Protoc. 2012;7(7):1299–1310.

40 Abrahams J, Campbell M, Packer N. Building a PGC-LC-MS N-glycan retention library and elution mapping resource. Glycoconjugate journal. 2018;35(1):15–29.

41 McCart Reed AE, Saunus JM, Ferguson K, Niland C, Simpson PT, Lakhani SR. The Brisbane Breast Bank. Open Journal of Bioresources. 2018;5:5.

42 Rivera-Franco MM, Leon-Rodriguez E. Delays in Breast Cancer Detection and Treatment in Developing Countries. Breast Cancer: Basic and Clinical Research. 2018;12:1–5.

43 Samraj AN, Pearce OM, Laubli H, et al. A red meat-derived glycan promotes inflammation and cancer progression. Proceedings of the National Academy of Sciences of the United States of America. 2015;112(2):542–547.

44 Tangvoranuntakul P, Gagneux P, Diaz S, et al. Human uptake and incorporation of an immunogenic nonhuman dietary sialic acid. Proceedings of the National Academy of Sciences of the United States of America. 2003;100(21):12045–12050.

45 Dhar C, Sasmal A, Varki A. From “Serum Sickness” to “Xenosialitis“: Past, Present, and Future Significance of the Non-human Sialic Acid Neu5Gc. Frontiers in immunology. 2019;10:807.

46 Bousquet PA, Sandvik JA, Jeppesen Edin NF, Krengel U. Hypothesis: Hypoxia induces de novo synthesis of NeuGc gangliosides in humans through CMAH domain substitute. Biochemical and biophysical research communications. 2018;495(1):1562–1566.

