## Supplementary Information for "*N*-glycolylneuraminic acid serum biomarker levels are elevated in breast cancer patients at all stages of disease"

##### Supplementary Methods

*Development of Glycoprotein Units (GPUs) standard curve for normalization of data from the SubB2M-$_{A12}$-SPR assays*

To generate an internal calibration curve, the Neu5Gc-containing glycoproteins bovine Alpha-1-acid glycoprotein (bAGP) (Sigma-Aldrich, Cat No. G3643) and human cancer antigen 125 (CA125) purified from a human ovarian carcinoma cell line (MyBioSource, San Diego, USA, Cat No. MBS318371) were combined at starting concentrations of 15 µg/ml and 15 units/ml, respectively, in 0.5 % normal human serum (equivalent to 3000 µg/ml bAGP and 3000 units/ml CA125 in 100 % serum). This glycoprotein mixture was two-fold serially diluted down to 14.65 ng/ml and 0.0146515 units/ml, respectively, in 0.5 % normal human serum. This concentration range of glycoprotein standards was run before every set of serum samples analyzed. RUs for each concentration of the GPU standard mixture were determined by subtracting binding due to SubB$_{A12}$ (flow cell 4) from binding due to SubB2M on flow cell 2 and flow cell 3. The RUs obtained for the highest concentration standard was considered 100 GPUs. The resulting standard curve was used to convert SPR RUs, taken at the point of stability in the generated sensorgram, to GPUs. The presence of Neu5Gc on both standard glycoproteins was confirmed by mass spectrometry as described previously (17) and below.

*Mass spectrometry glycomic analysis of standard glycoproteins*

Glycoproteins (10 µg bAGP and 116.83 units CA125) were immobilised onto PVDF membrane (Millipore) and *N*-glycans released by overnight incubation with 0.5 µl PNGase F (New England BioLabs 500,000 units/ml) in 10 µl water at 37 °C. Released *N*-glycans were reduced to alditols with 0.5 M NaBH4 in 50 mM KOH for 3 h at 50 °C. The reduction was quenched with 1 µl glacial acetic acid desalted using AG50W-X8 cation exchange resin.

*O*-glycans were released from PNGase F treated proteins by reductive β-elimination. PVDF spots were incubated in 20 µL of 0.5 M $NaBH_4$ in 50 mM KOH at 50 °C for 16 hours and desalted as described for the *N*-glycans.

PGC-LC-ESI-MS *N*-glycans were analysed using a Hypercarb PGC column (3 µm, 100 mm × 180 µm, Thermo Scientific). *N*-glycans were separated over a 90 min and *O*-glycans over a 60 min gradient of 1–90 % of acetonitrile in 10 mM ammonium bicarbonate (vol/vol) at a flow rate of 1 µL/min using a Dionex ultimate HPLC (Thermo Scientific) interfaced with an amaZon Speed ESI-IT mass spectrometer (Bruker Bruker Daltonics, Germany).

The MS spectra were acquired in negative ion mode over a mass range of 450 to 2200 m/z. The following MS settings were used: drying gas temperature: 180 °C, drying gas flow: 5 L/min, nebulizer gas: 9 psi, capillary 3400 V. Ions were detected in ion charge control (ICC) (target: 50,000 ions) with an accumulation time of 200 ms. Induced collision was performed at 35 % normalised collision energy and an isolation window of 4 m/z). Instrument control, data acquisition and processing were performed with Bruker DataAnalysis software version 4.2 (Bruker Daltonics, Germany).

#### Supplementary Figures

**Figure S1. Glycan array analysis of SubB2M and $SubB_{A12}$ using a Z-Biotech Neu5Ac/Neu5Gc array. A)** Glycan array result of SubB2M and $SubB_{A12}$ performed using the Z-Biotech Neu5Gc/Neu5Ac N-Glycan Array. Histogram represents the average relative fluorescent units of binding to each of the numbered structures shown in **B**. For structure ID see http://www.zbiotech.com/neu5gc-xenoantigen-microarray.html and http://nebula.wsimg.com/deda6829116ce09edb871bd7ce7cde6c?AccessKeyId=B5CD53DB37409833427C&disposition=0&alloworigin=1 for further information.

**Figure S2. Characterization of human CA125 *O*-glycosylation and bovine Alpha-1-acid glycoprotein (bAGP) by PGC-LC-MS/MS.** Annotated Base Peak Chromatogram of the total **A)** *O*-glycome released from CA125 and Extracted ion chromatogram of m/z $681.32^-$ (Neu5Gc) and $665.32^-$ (Neu5Ac) and **B)** *N*-glycome released from bAGP and Extracted ion chromatogram of m/z $1127.4^{2-}$ (Neu5Gc) and $1111.4^{2-}$ (Neu5Ac). Confirmation of **C)** Neu5Gc (m/z $681.32^-$) and Neu5Ac (m/z $665.32^-$) containing *O*-glycan structures by MS/MS fragmentation and **D**) Neu5Gc (m/z $1127.4^{2-}$) and Neu5Ac (m/z $1111.4^{2-}$) containing glycan structures by MS/MS fragmentation.

**Figure S3. A representative Glycoprotein Units (GPUs) standard curve.** Bovine AGP (MW = 41-43 kDa; ~50 %/50 % Neu5Ac/Neu5Gc; high total sialic acids) and human CA125 (MW = >200 kDa, 5-10 % Neu5Gc; low total sialic acid) were combined at starting concentrations of 15 µg/ml and 15 units/ml, respectively, in 0.5 % normal human serum. This glycoprotein mixture was two-fold serially diluted down to 14.65 ng/ml and 0.0146515 units/ml, respectively, in 0.5 % normal human serum. The Response Units (RUs) for each concentration of the standard mixture were determined by subtracting binding due to SubB$_{A12}$ (flow cell 4) from binding due to SubB2M on flow cell 2 and flow cell 3. RUs obtained for the highest concentration standard was considered 100 GPUs. FC2 = flow cell 2; FC3 = flow cell 3.

**Figure S4. ROC curves depicting the ability of serum Neu5Gc levels determined by the optimized SubB2M-$_{A12}$-SPR assay to distinguish Stage I – IV ovarian cancer patients from cancer-free (normal) individuals.** Sensitivity% (true positive rate; ability to detect disease) is plotted against 100 %-specificity% (false positive rate or 100 %-true negative rate; ability to detect lack of disease). ROC analyses were performed with the data shown in Figure 2B using Graphpad Prism 8.0

**Figure S5. ROC curves depicting the ability of serum Neu5Gc levels to distinguish Stage I – IV breast cancer patients from normal (cancer-free) individuals.** Sensitivity% (true positive rate; ability to detect disease) is plotted against 100 %-specificity% (false positive rate or 100 %-true negative rate; ability to detect lack of disease). ROC analyses were performed with the data shown in Figure 3 using Graphpad Prism 8.0

**Figure S6. Serum Neu5Gc levels determined by SubB2M-$_{A12}$-SPR assay for A) relapse cases and B) remission cases from the Circ.BR cohort.** The mean GPUs from duplicate analyses for each serum sample are shown.

Error bars = ± 1 SD from the mean for each group. Two independent assays were performed with both showing the same trends. Results from one assay are shown. Clinical information for each patient is shown in the top right of each plot with treatment history and metastases overlaid. ALND: Axillary lymph node dissection, ILC: Invasive Lobular Carcinoma Mast: mastectomy, SNB: sentinel node biopsy, WLE: wide local excision, XRT: radiation therapy. Detailed information for each patient in the Circ.BR cohort are shown in Supplementary Table 3.

**Supplementary Table 4. Optimal cut-off values, sensitivity and specificity for distinguishing Stage I, II, III and IV ovarian cancer patients from normal (cancer-free) individuals using serum Neu5Gc levels determined by optimized SubB2M-SPR assay before and after $SubB_{A12}$ subtraction.** Sensitivity and specificity were determined from the Receiver operating characteristic (ROC) curves (**Figure S4**). Optimal cut-off values were selected to give the maximum sum of sensitivity and specificity.

| | **Before $SubB_{A12}$ subtraction** | **After $SubB_{A12}$ subtraction** |
|---|---|---|
| **Normal vs Stage I** | >9.02 GPUs (91.67 % sensitivity, 100 % specificity) | >6.90 GPUs (100 % sensitivity, 100 % specificity) |

| | | |
|---|---|---|
| **Normal vs Stage II** | >8.83 GPUs (90.91 % sensitivity, 94.45 % specificity) | >6.88 GPUs (100 % sensitivity, 100 % specificity) |
| **Normal vs Stage III** | >14.50 GPUs (100 % sensitivity, 100 % specificity) | >16.40 GPUs (100 % sensitivity, 100 % specificity) |
| **Normal vs Stage IV** | >10.49 GPUs (100 % sensitivity, 100 % specificity) | >11.87 GPUs (100 % sensitivity, 100 % specificity) |

**Supplementary Table 5. Optimal cut-off and area under the curve (AUC) values for distinguishing Stage I, II, III and IV breast cancer patients from normal (cancer-free) individuals using serum Neu5Gc levels.** Sensitivity and specificity were determined from the receiver operating characteristic (ROC) curves (**Figure S5**). Optimal cut-off values were selected to give the maximum sum of sensitivity and specificity.

| | **Optimal cut-off** | **ROC AUC** |
|---|---|---|
| **Normal vs Stage I** | >10.55 GPU (sensitivity = 95.83 %, specificity = 100 %) | 0.9583 |
| **Normal vs Stage II** | >10.49 GPU (sensitivity = 100 %, specificity = 100 %) | 1.000 |
| **Normal vs Stage III** | >14.54 GPU (sensitivity = 100 %, specificity = 100 %) | 1.000 |

| | | |
|---|---|---|
| **Normal vs Stage IV** | >19.87GPU (sensitivity = 100%, specificity = 100%) | 1.000 |

**Supplementary Table S1.** Supplementary glycan microarray document based on MIRAGE guidelines DOI: 10.1093/glycob/cww118.

| Classification | Guidelines |
|---|---|
| **1. Sample: Glycan Binding Sample** | |
| Description of Sample | Sample names:<br>*Escherichia coli* SubB2M and SubB$_{A12}$.<br><br>Origin: B subunit pentameric toxins produced as a recombinant protein in *E. coli*.<br><br>Method of preparation:<br>The preparation of SubB2M and SubB$_{A12}$ are explained in the Materials and Methods section. |
| Sample modifications | SubB2M and SubB$_{A12}$ are hexahistidine-tagged proteins. |
| Assay protocol | Please see Materials and Methods and appended manufacture's manual. |
| **2. Glycan Library** | |
| Glycan description for defined glycans | Arrays used are the Z-Biotech Neu5Gc/Neu5Ac N-Glycan Array. Glycans in this study are listed in Supplementary Figure S1 and are outlined below. |
| Glycan description for undefined glycans | N/A. |
| Glycan modifications | N/A |
| **3. Printing Surface; e.g., Microarray Slide** | |
| Description of surface | NHS matrix slides |
| Manufacturer | Schott Nexterion |
| Custom preparation of surface | N/A. |
| Non-covalent Immobilisation | N/A. |
| **4. Arrayer (Printer)** | |

| | |
|---|---|
| Description of Arrayer | See Z-Biotech |
| Dispensing mechanism | See Z-Biotech |
| Glycan deposition | See Z-Biotech |
| Printing conditions | See Z-Biotech |
| **5. Glycan Microarray with "Map"** | |
| Array layout | See page 3. |
| Glycan identification and quality control | Arrays are quality controlled as described on page 6. |
| **6. Detector and Data Processing** | |
| Scanning hardware | Innopsys InnoScan 1100AL (Lasers: 488 nM, 532 nM with two filter sets for analysis at 532 and 595 nM), 635 nM) scanner. |
| Scanner settings | Scanning resolution: 10 µM<br>Laser channel: 532 nM operating 532 nM excitation filter set.<br>PMT: 20 % gain<br>Scan powers: Low laser power. |
| Image analysis software | Innopsys MAPIX. |
| Data processing | Data was exported as a CSV file and exported to Microsoft Excel. |
| **7. Glycan Microarray Data Presentation** | |
| Data presentation | Data is presented as histograms in Figure S1. |
| **8. Interpretation and Conclusion from Microarray Data** | |
| Data interpretation | We only use glycan arrays as a yes/no binding tool. Due to this we look only at binding that is unambiguously above background vs lack of binding above background. |
| Conclusions | SubB2M is specific for Neu5Gc, $SubB_{A12}$ does not bind to any sialic acid containing glycans. |

Print Layout

16-subarray Slide

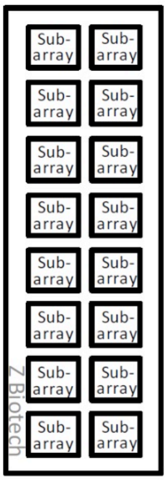

**Array Map:**

| | | | | | | | | | | | | | | | |
|---|---|---|---|---|---|---|---|---|---|---|---|---|---|---|---|
| GC-1 | GC-1 | GC-1 | GC-2 | GC-2 | GC-2 | GC-3 | GC-3 | GC-3 | GC-4 | GC-4 | GC-4 | GC-5 | GC-5 | GC-5 | NC1 |
| GC-6 | GC-6 | GC-6 | GC-7 | GC-7 | GC-7 | GC-8 | GC-8 | GC-8 | GC-9 | GC-9 | GC-9 | GC-10 | GC-10 | GC-10 | NC1 |
| GC-11 | GC-11 | GC-11 | GC-12 | GC-12 | GC-12 | GC-13 | GC-13 | GC-13 | GC-14 | GC-14 | GC-14 | GC-15 | GC-15 | GC-15 | NC1 |
| GC-16 | GC-16 | GC-16 | GC-17 | GC-17 | GC-17 | GC-18 | GC-18 | GC-18 | GC-19 | GC-19 | GC-19 | GC-20 | GC-20 | GC-20 | PC1 |
| GC-21 | GC-21 | GC-21 | GC-22 | GC-22 | GC-22 | GC-23 | GC-23 | GC-23 | GC-24 | GC-24 | GC-24 | GC-25 | GC-25 | GC-25 | PC1 |
| GC-26 | GC-26 | GC-26 | GC-27 | GC-27 | GC-27 | GC-28 | GC-28 | GC-28 | GC-29 | GC-29 | GC-29 | GC-30 | GC-30 | GC-30 | PC1 |
| GC-31 | GC-31 | GC-31 | GC-32 | GC-32 | GC-32 | GC-33 | GC-33 | GC-33 | GC-34 | GC-34 | GC-34 | GC-35 | GC-35 | GC-35 | PC2 |
| GC-36 | GC-36 | GC-36 | GC-37 | GC-37 | GC-37 | GC-38 | GC-38 | GC-38 | GC-39 | GC-39 | GC-39 | GC-40 | GC-40 | GC-40 | PC2 |
| AC-1 | AC-1 | AC-1 | AC-2 | AC-2 | AC-2 | AC-3 | AC-3 | AC-3 | AC-4 | AC-4 | AC-4 | AC-5 | AC-5 | AC-5 | PC2 |
| AC-6 | AC-6 | AC-6 | AC-7 | AC-7 | AC-7 | AC-8 | AC-8 | AC-8 | AC-9 | AC-9 | AC-9 | AC-10 | AC-10 | AC-10 | PC3 |
| AC-11 | AC-11 | AC-11 | AC-12 | AC-12 | AC-12 | AC-13 | AC-13 | AC-13 | AC-14 | AC-14 | AC-14 | AC-15 | AC-15 | AC-15 | PC3 |
| AC-16 | AC-16 | AC-16 | AC-17 | AC-17 | AC-17 | AC-18 | AC-18 | AC-18 | AC-19 | AC-19 | AC-19 | AC-20 | AC-20 | AC-20 | PC3 |
| AC-21 | AC-21 | AC-21 | AC-22 | AC-22 | AC-22 | AC-23 | AC-23 | AC-23 | AC-24 | AC-24 | AC-24 | AC-25 | AC-25 | AC-25 | PC4 |
| AC-26 | AC-26 | AC-26 | AC-27 | AC-27 | AC-27 | AC-28 | AC-28 | AC-28 | AC-29 | AC-29 | AC-29 | AC-30 | AC-30 | AC-30 | PC4 |
| AC-31 | AC-31 | AC-31 | AC-32 | AC-32 | AC-32 | AC-33 | AC-33 | AC-33 | AC-34 | AC-34 | AC-34 | AC-35 | AC-35 | AC-35 | PC4 |
| AC-36 | AC-36 | AC-36 | AC-37 | AC-37 | AC-37 | AC-38 | AC-38 | AC-38 | AC-39 | AC-39 | AC-39 | GC-41 | GC-41 | GC-41 | Marker |

Glycan list:

#### N-Glycan Identification List:

| Gc Glycan ID | Neu5Gc Glycans | Ac Glycan ID | Neu5Ac Glycans |
|---|---|---|---|
| GC-1 | N002G | AC-1 | N002 |
| GC-2 | N003G | AC-2 | N003 |
| GC-3 | N005G | AC-3 | N005 |
| GC-4 | N012G | AC-4 | N012 |
| GC-5 | N013G | AC-5 | N013 |
| GC-6 | N015G | AC-6 | N015 |
| GC-7 | N022G | AC-7 | N022 |
| GC-8 | N023G | AC-8 | N023 |
| GC-9 | N025G | AC-9 | N025 |
| GC-10 | N032G | AC-10 | N032 |
| GC-11 | N033G | AC-11 | N033 |
| GC-12 | N042G | AC-12 | N042 |
| GC-13 | N043G | AC-13 | N043 |
| GC-14 | N045G | AC-14 | N045 |
| GC-15 | N052G | AC-15 | N052 |
| GC-16 | N053G | AC-16 | N053 |
| GC-17 | N055G | AC-17 | N055 |
| GC-18 | N112G | AC-18 | N112 |
| GC-19 | N113G | AC-19 | N113 |
| GC-20 | N115G | AC-20 | N115 |
| GC-21 | N122G | AC-21 | N122 |
| GC-22 | N123G | AC-22 | N123 |
| GC-23 | N125G | AC-23 | N125 |
| GC-24 | N133G | AC-24 | N133 |
| GC-25 | N134G | AC-25 | N134 |
| GC-26 | N135G | AC-26 | N135 |
| GC-27 | N144G | AC-27 | N144 |
| GC-28 | N145G | | |
| GC-29 | N155G | AC-29 | N155 |
| GC-30 | N212G | AC-30 | N212 |
| GC-31 | N213G | AC-31 | N213 |
| GC-32 | N215G | AC-32 | N215 |
| GC-33 | N222G | AC-33 | N222 |
| GC-34 | N223G | AC-34 | N223 |
| GC-35 | N225G | AC-35 | N225 |
| GC-36 | N233G | AC-36 | N233 |
| GC-37 | N235G | | |
| GC-38 | N245G | | |
| GC-39 | N255G | AC-39 | N255 |
| GC-40 | N003G1 | | |
| GC-41 | N003G2 | | |

### Neu5Gc and Neu5Ac N-Glycans

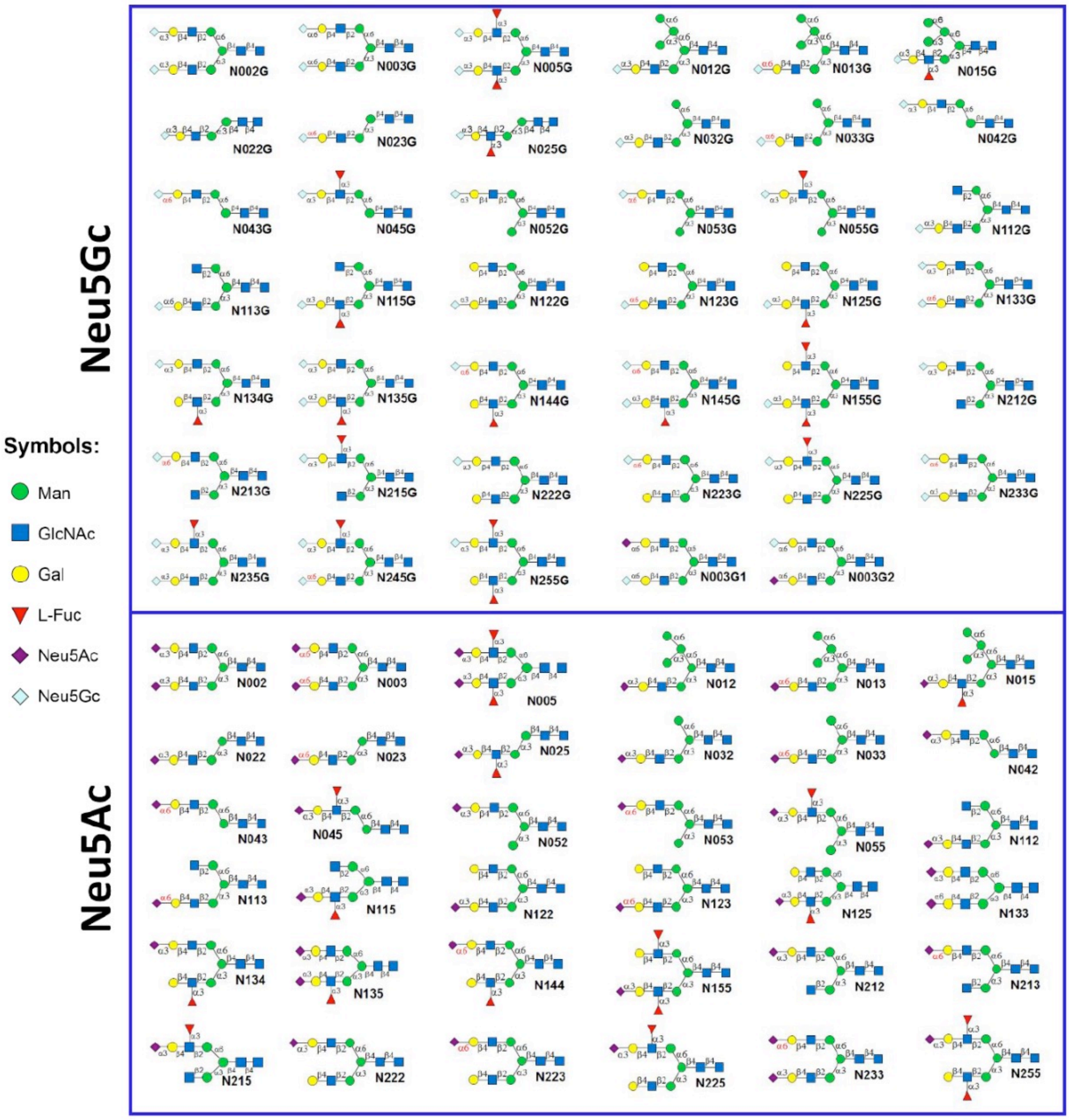

QC

Example 1: Neu5Gc/Neu5Ac array on 8 subarray format. A subarray assayed with a biotinylated SNA target (20 µg/ml), followed by streptavidin-Cy3 (1 µg/ml). The array was scanned with GenePix scanner at 500 PMT and 100% laser power at 532 nm wavelength. The positive control shows binding as expected. N-glycans containing α-2,6 Neu5Gc and α-2,6 Neu5Ac show binding as expected. Analysis of the fluorescence intensity reveals that Neu5Gc-sialylated glycans bind more strongly.

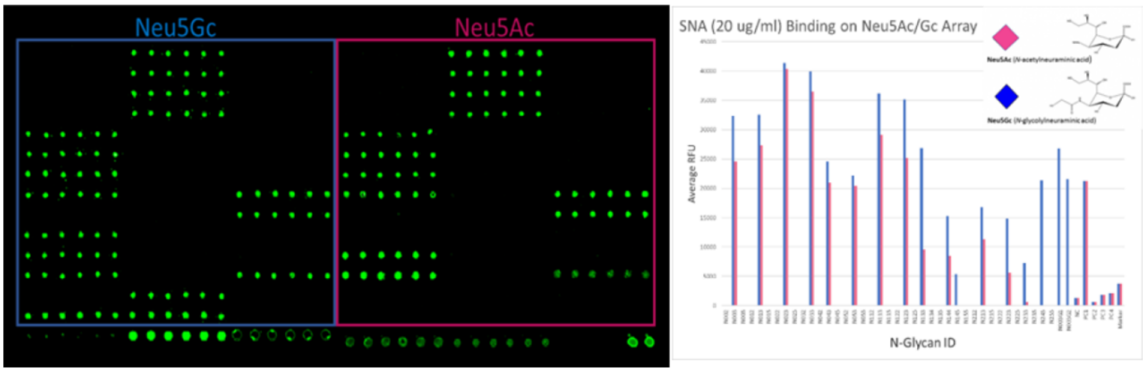

Example 2: Neu5Gc/Neu5Ac array on 8 subarray format. A subarray assayed with a biotinylated WGA target (10 µg/ml), followed by streptavidin-Cy3 (1 µg/ml). The array was scanned with GenePix scanner at 450 PMT and 100% laser power at 532 nm wavelength. The positive control shows binding as expected. Most N-glycans show binding as expected. Analysis of the fluorescence intensity reveals that Neu5Ac-sialylated glycans bind more strongly.

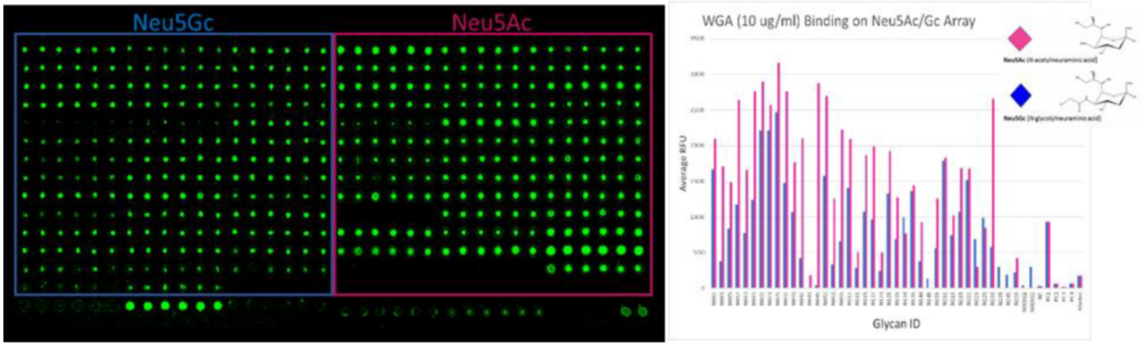

**Supplementary Table S2. Details for each of the normal (cancer-free) individuals and breast cancer patients used in this study.** Patient details were provided by the Victorian Cancer Biobank with informed written consent from each subject. Abbreviations: ALND: axillary lymph node dissection; DCIS: ductal carcinoma in-situ; G1: Grade 1; G2: Grade 2; G3: Grade 3; LCIS: lobular carcinoma in-situ; LVSI: lymph vascular space invasion; NST: no special type. Information on vital status and/or recurrence was not available from the VCB for all patients.

| Specimen no. | Age | Breast cancer stage | Type of breast cancer | Vital status | Date of death | Survival time (years)* | Recurrence | Neu5Gc levels (GPUs) |
|---|---|---|---|---|---|---|---|---|
| 07AH130 | 50 | N/A | Cancer-free | | | | | 6.083 |
| 08AH714 | 42 | N/A | Cancer-free | | | | | 6.326 |
| 09AH320 | 47 | N/A | Cancer-free | | | | | 4.972 |
| 09AH434 | 42 | N/A | Cancer-free | | | | | 4.278 |
| 09AH794 | 57 | N/A | Cancer-free | | | | | 10.077 |
| 09AH796 | 52 | N/A | Cancer-free | | | | | 8.653 |
| 09AH820 | 38 | N/A | Cancer-free | | | | | 5.076 |
| 12EH0028 | 53 | N/A | Cancer-free | | | | | 4.694 |
| 12EH0114 | 72 | N/A | Cancer-free | | | | | 5.563 |
| 15EH0238 | 51 | N/A | Cancer-free | | | | | 7.785 |
| 15EH0234 | 79 | N/A | Cancer-free | | | | | 7.125 |
| 15EH0228 | 62 | N/A | Cancer-free | | | | | 6.257 |
| 16EH0287 | 61 | N/A | Cancer-free | | | | | 2.472 |
| 16EH0217 | 76 | N/A | Cancer-free | | | | | 5.806 |
| 16EH0396 | 74 | N/A | Cancer-free | | | | | 3.965 |
| 16EH0397 | 50 | N/A | Cancer-free | | | | | 0.423 |
| 17EH0084 | 54 | N/A | Cancer-free | | | | | 4.556 |
| 17EH0260 | 46 | N/A | Cancer-free | | | | | 7.299 |
| 17EH0268 | 63 | N/A | Cancer-free | | | | | 5.528 |
| 17EH0314 | 46 | N/A | Cancer-free | | | | | 3.792 |
| 17EH0349 | 93 | N/A | Cancer-free | | | | | 3.792 |
| 17EH0211 | 52 | N/A | Cancer-free | | | | | 4.937 |
| 02PM1046 | 80 | I | Invasive lobular carcinoma (G2) | | | | | 20.702 |

| | | | | | | | | |
|---|---|---|---|---|---|---|---|---|
| 04PM1332 | 55 | I | Invasive ductal carcinoma (G2) with associated DCIS (high grade) | A | N/A | | | 37.856 |
| 09PM0065 | 53 | I | Invasive ductal carcinoma (G3) with associated DCIS (high grade) | A | N/A | | | 24.904 |
| 15PM0686 | 38 | I | Invasive ductal carcinoma (G2) with associated DCIS (high grade) | A | N/A | | | 11.014 |
| 15PM0997 | 59 | I | Invasive mixed micropapillary and ductal/NST carcinoma (G3) and DCIS (high grade) | | | | | 24.349 |
| 11MH0317 | 43 | I | Invasive ductal carcinoma NST (G3) | A | N/A | | | 25.460 |
| 12MH0314 | 44 | I | Invasive ductal carcinoma NST (G2) with associated DCIS (high grade) | A | N/A | | | 25.251 |
| 11MH0052 | 68 | I | Invasive ductal carcinoma (G2) with associated DCIS (low grade) | | | | | 16.188 |
| 11MH0554 | 61 | I | Invasive ductal carcinoma NST (G2) with associated DCIS (high grade) | A | N/A | | | 22.508 |
| 13MH1003 | 66 | I | Invasive ductal carcinoma (G1) with associated DCIS (low grade) | | | | | 17.299 |
| 10RMH737 | 58 | I | Invasive ductal carcinoma NST (G2) with associated DCIS (high grade) | | | | | 14.799 |
| 11MH0567 | 47 | I | Invasive ductal carcinoma with lobular features (G2) with associated DCIS (low grade) | | | | | 12.369 |
| 09RMH083 | 60 | I | Invasive ductal carcinoma NST (G2) with extensive DCIS (high grade) | | | | | 35.634 |
| 10RMH281 | 54 | I | Invasive ductal carcinoma NST (G3) | A | N/A | | | 22.717 |
| 10RMH383 | 77 | I | Invasive mucinous adenocarcinoma (G2) | | | | | 30.391 |
| 10RMH514 | 81 | I | Invasive lobular carcinoma NST (G2) with associated LCIS | A | N/A | | | 35.947 |
| 12MH0215 | 35 | I | Invasive ductal carcinoma NST (G2) | | | | | 14.869 |
| 12MH0481 | 56 | I | Invasive ductal carcinoma NST (G2) with associated DCIS (high grade) | | | | | 23.133 |
| 12MH1013 | 65 | I | Invasive ductal adenocarcinoma NST (G1) with DCIS (low grade) | A | N/A | | | -3.674 |
| 12MH1291 | 56 | I | Invasive ductal adenocarcinoma NST (G2) | A | N/A | | | 26.849 |

| | | | | | | | | |
|---|---|---|---|---|---|---|---|---|
| 13MH1223 | 57 | I | Invasive ductal carcinoma NST (G2) with DCIS (intermediate grade) | | | | | 24.210 |
| 14MH0232 | 51 | I | Invasive ductal carcinoma NST (G1) | A | N/A | | | 22.091 |
| 14MH0240 | 43 | I | Invasive ductal carcinoma NST (G1) with LCIS (moderate) | A | N/A | | | 28.446 |
| 16MH1550 | 48 | I | Lobular carcinoma NST (G3) with associated LCIS | A | N/A | | | 20.772 |
| 02PM0520 | 58 | II | Invasive ductal carcinoma NST (G3) with associated DCIS (high grade) & metastatic adenocarcinoma | A | N/A | | | 23.967 |
| 04PM0880 | 45 | II | Invasive ductal carcinoma (G3) with associated DCIS (minor high grade) & metastatic carcinoma | A | N/A | | | 17.543 |
| 05PM1100 | 31 | II | Mixed invasive ductal carcinoma (NTS and mucinous) (G2) with associated DCIS (high grade) & metastatic tumor of sentinel nodes | A | N/A | | | 11.605 |
| 05PM1168 | 55 | II | Invasive ductal carcinoma (G3) with associated DCIS (high grade) & metastatic carcinoma | D | 30/05/2008 | 3 | | 13.410 |
| 06PM0542 | 26 | II | Invasive mucinous carcinoma (G1 & G2) with associated DCIS (high grade) | | | | | 11.257 |
| 08PM1308 | 59 | II | Invasive ductal carcinoma NST (G3) & metastatic carcinoma | A | N/A | | | 17.786 |
| 08PM1949 | 49 | II | Invasive lobular carcinoma (G2) (post chemotherapy) & metastatic carcinoma | A | N/A | | | 15.772 |
| 15PM0781 | 44 | II | Invasive ductal carcinoma NST (G3) & metastatic carcinoma | A | N/A | | | 10.910 |
| 14PM0388 | 67 | II | Invasive ductal carcinoma NST (G3) with associated DCIS (high grade) | | | | | 12.542 |
| 15PM1021 | 53 | II | Invasive ductal carcinoma NST (G2) with associated DCIS (intermediate to high grade) | A | N/A | | | 10.945 |
| 15PM1119 | 51 | II | Mixed invasive carcinoma (micropapiliary and NST) (G3) with associated DCIS (low and high grade) & metastatic carcinoma | A | N/A | | | 18.619 |
| 13MH0429 | 51 | II | Invasive ductal carcinoma (G2) with associated DCIS (intermediate grade) & metastatic carcinoma | D | 26/04/2017 | 4.08 | | 13.063 |
| 16MH1592 | 47 | II | Invasive ductal carcinoma NST (G3) with minor DCIS | A | N/A | | | 28.585 |

| | | | | | | | | |
|---|---|---|---|---|---|---|---|---|
| 01PM0361 | 75 | II | Invasive ductal carcinoma NST (G3) with DCIS (high grade) | D | 03/01/2003 | 1.33 | | 20.286 |
| 01PM0632 | 71 | II | Invasive ductal carcinoma NST (G3) | D | 06/06/2003 | 1.75 | No | 23.411 |
| 03PM0734 | 47 | II | Invasive ductal carcinoma NST (G2) & metastatic carcinoma | | | | | 22.091 |
| 05PM2377 | 49 | II | Invasive ductal carcinoma NST (G1) with DCIS (low grade) | A | N/A | | Yes | 30.738 |
| 08PM1773 | 79 | II | Invasive ductal carcinoma NST (G3) with DCIS (high grade) | | | | | 20.598 |
| 08PM1785 | 54 | II | Invasive ductal carcinoma NST (G3) with DCIS (high grade) & metastatic carcinoma | A | N/A | | | 20.771 |
| 09PM0040 | 51 | II | Invasive ductal carcinoma NST (G1) with DCIS (high grade) | A | N/A | | Yes | 16.154 |
| 10PM0659 | 31 | II | Invasive ductal carcinoma (G2) with neuroendocrine differentiation | | | | | 27.092 |
| 10PM2156 | 78 | II | Invasive ductal carcinoma NST (G3) with DCIS (high grade) | D | 18/03/2017 | 6.33 | No | 17.022 |
| 12PM0321 | 46 | II | Invasive ductal carcinoma NST (G1) with DCIS (low to intermediate grade) & metastatic carcinoma | A | N/A | | | 15.945 |
| 13PM0443 | 39 | II | Invasive ductal (NST) (G3) and lobular carcinoma with DCIS (intermediate to high grade) | D | 20/02/2015 | 1.83 | Yes | 16.223 |
| 01PM0503 | 49 | III | Invasive ductal carcinoma (G3) with associated DCIS (high grade) of right breast | D | 09/03/2002 | 0.42 | | 65.254 |
| 03PM0187 | 57 | III | Invasive ductal carcinoma NTS (G3) with associated DCIS (high grade) of left breast with invasion of skeletal muscle and lymphatic channels | | | | | 30.530 |
| 03PM1038 | 32 | III | Invasive ductal carcinoma (G3) with associated DCIS (high grade) of right breast & metastatic carcinoma | D | 18/12/2004 | 1.42 | | 31.016 |
| 04PM0931 | 51 | III | Invasive ductal carcinoma NST (G3) with possible DCIS of right breast & metastatic carcinoma, advanced left breast cancer | D | 06/01/2005 | 3 | | 55.879 |
| 08PM1472 | 57 | III | Invasive ductal carcinoma (G3) with associated DCIS (high grade) of right breast & metastatic carcinoma | | | | | 28.967 |
| 09PM1880 | 55 | III | Invasive ductal carcinoma (G3) (basal type differentiation) with associated DCIS (high grade) of | A | N/A | | | 41.572 |

| | | | | | | | | | |
|---|---|---|---|---|---|---|---|---|---|
| | | | left breast & metastatic carcinoma | | | | | | |
| 11PM0575 | 33 | III | Invasive mucinous carcinoma (G1) of left breast & metastatic carcinoma | A | N/A | | | | 33.724 |
| 11PM1136 | 42 | III | Invasive ductal carcinoma (G3) with associated DCIS (high grade) of right breast & metastatic carcinoma | D | 31/08/2013 | 2.17 | | | 37.683 |
| 14PM1036 | 83 | III | Invasive ductal carcinoma NST (G3) with associated DCIS (high grade) of right breast & metastatic carcinoma | A | N/A | | | | 32.856 |
| 15PM0518 | 53 | III | Invasive ductal adenocarcinoma (G3) with associated LCIS (high grade) of left breast & metastatic carcinoma | A | N/A | | | | 52.059 |
| 15PM0965 | 56 | III | Invasive carcinoma (basal) (G3) with associated DCIS (high grade) of left breast & metastatic carcinoma | A | N/A | | | | 73.658 |
| 11MH0137 | 43 | III | Invasive ductal carcinoma NST (G3) with associated DCIS (high grade) of right breast & metastatic carcinoma | A | N/A | | | | 30.078 |
| 09RMH727 | 45 | III | Poorly differentiated invasive ductal carcinoma (G3) & metastatic carcinoma | A | N/A | | | | 53.378 |
| 10RMH275 | 69 | III | Invasive ductal carcinoma NTS (G1) with DCIS (intermediate to high grade) & metastatic carcinoma | | | | | | 42.857 |
| 12MH0211 | 48 | III | Invasive ductal carcinoma (G2) with metastatic ductal carcinoma | A | N/A | | | | 41.259 |
| 12MH0483 | 56 | III | Micropapillary invasive ductal carcinoma (G3) | A | N/A | | | Yes | 44.142 |
| 13MH0077 | 56 | III | Invasive lobular carcinoma (G1) & metastatic lobular carcinoma | A | N/A | | | | 53.031 |
| 18MH0981 | 36 | III | Invasive ductal carcinoma NTS (G3) with DCIS (low and high grade) | A | N/A | | | | 26.363 |
| 01PM0213 BLD | 66 | III | Invasive ductal carcinoma (moderately differentiated) (G2/3) with DICS (intermediate) & metastatic carcinoma | | | | | | 31.120 |
| 01PM0463 | 81 | III | Invasive ductal carcinoma (G2) & metastatic carcinoma | D | 12/09/2003 | 2 | | Yes | 38.968 |
| 01PM0443 | 76 | III | Invasive ductal carcinoma (G3) with some lobular pattern & metastatic carcinoma | D | 20/11/2002 | 1.17 | | | 19.001 |

| | | | | | | | | |
|---|---|---|---|---|---|---|---|---|
| 02PM1143 | 39 | III | Invasive ductal carcinoma (G2) with associated DCIS (high grade) & metastatic carcinoma | D | 05/01/2011 | 8.08 | | 26.883 |
| 03PM0227 BLD | 57 | III | Invasive ductal carcinoma NTS (G3) with DCIS (high grade) | | | | | 40.808 |
| 08PM2010 BLD | 50 | III | Invasive ductal carcinoma (G3) | A | N/A | | Yes | 27.890 |
| 17MH0909bl | 49 | IV | Invasive ductal carcinoma (G3) with associated DCIS (high grade) of left breast with foci of LVSI & metastatic carcinoma | D | 06/05/2018 | 0.83 | | 181.200 |
| 17MH0907bl | 76 | IV | Metastatic breast carcinoma (stable stage IV for 7 years, primary lung cancer) | | | | | 203.389 |
| 15MH1848bl | 54 | IV | Metastatic adenoid cystic carcinoma of right breast (G3) (also adenoid cystic carcinoma of brain consistent with primary breast cancer) | | | | | 134.566 |
| 15MH1826bl | 45 | IV | Invasive adenocarcinoma NST (right breast, G3; left breast, G2) with associated DCIS (high grade) & metastatic carcinoma (clavicular head; 6/7 right lymph nodes; 7/7 left lymph nodes), history of Non-Hodgkin lymphoma | A | N/A | | | 138.211 |
| 15MH1733bl | 27 | IV | Invasive ductal carcinoma (G3) with associated DCIS (high grade) of left breast & metastatic carcinoma (4/5 sentinel nodes; 10/11 lymph nodes with extravascular spread) | A | N/A | | | 92.548 |
| 13MH1140bl | 62 | IV | Invasive lobular carcinoma of left breast (G2); Invasive lobular carcinoma (G2) and invasive ductal carcinoma NST (G3) of right breast & metastatic carcinoma (3/18 right axillary lymph nodes) | A | N/A | | | 74.595 |
| 13MH0914bl | 42 | IV | Invasive ductal carcinoma (G3) of left breast; recurrent disease; metastasis to the vertebrae | D | 07/12/2013 | 0.25 | Yes | 59.108 |
| 13MH0767bl | 48 | IV | Invasive ductal carcinoma (G2) of left breast & metastatic carcinoma (bone metastasis; 1/1 lymph nodes) | D | 03/02/2016 | 2.5 | | 62.962 |
| 13MH0529bl | 62 | IV | Invasive ductal adenocarcinoma (G3) of right breast & metastatic carcinoma (pulmonary metastasis, 2/5 lymph nodes) | D | 02/03/2016 | 2.75 | | 50.774 |

| | | | | | | | | | |
|---|---|---|---|---|---|---|---|---|---|
| 13MH0217bl | 61 | IV | Invasive ductal adenocarcinoma (G3) of right breast & metastatic adenocarcinoma (9/12 lymph nodes with extravascular extension; bone metastasis) | A | N/A | | | | 36.572 |
| 12MH1318 | 41 | IV | Invasive ductal carcinoma (G2) of left breast with associated DCIS (high grade) & metastatic carcinoma (1/13 lymph nodes; pulmonary metastasis) | A | N/A | | | | 67.650 |
| 12MH0226 | 66 | IV | Invasive ductal carcinoma (G3) of right breast & metastatic carcinoma (19/21 lymph nodes; pulmonary metastasis) | D | 12/12/2016 | 3.75 | | | 55.635 |
| 02PM0246 | 73 | IV | Longstanding metastatic breast carcinoma (bone, pulmonary and adrenal metastasis) | | | | | | 176.060 |
| 02PM0320 | 64 | IV | Invasive primary breast adenocarcinoma (uterus, ovary, fallopian tube and omentum metastasis) | | | | | | 130.884 |
| 05PM1349 | 48 | IV | Invasive lobular carcinoma (G2) with minor mucinous component & DCIS (intermediate and high grade) of left breast | D | 28/09/2005 | 0.25 | | | 126.856 |
| 06PM0159 | 54 | IV | Primary breast carcinoma & metastatic carcinoma (oesophagus) | D | 26/12/2006 | 0.83 | | | 88.416 |
| 08PM1819 | 57 | IV | Invasive ductal carcinoma of breast (metastasis to left shoulder/chest wall) | | | | | | 77.338 |
| 08PM1958 | 47 | IV | Metastatic primary breast carcinoma (left femoral head) | | | | | | 39.002 |
| 10PM1401 | 60 | IV | Metastatic primary breast adenocarcinoma (metastasis to brain and spine) | | | | | | 62.476 |
| 11PM0548 | 51 | IV | Metastatic primary breast carcinoma (metastasis to brain) | | | | | | 46.607 |
| 11PM1336 | 39 | IV | Metastatic breast carcinoma (adenocarcinoma of ovary and fallopian tubes) | A | N/A | | | | 29.661 |
| 11PM1339 | 39 | IV | Invasive ductal carcinoma NST (G3) with associated DCIS (intermediate to high grade) of both breasts & metastatic breast adenocarcinoma (metastasis to ovaries) | D | 29/06/2014 | 2.83 | | | 53.760 |
| 13PM0154 | 72 | IV | Metastatic lobular carcinoma (15/18 lymph nodes of left axillary; 10/36 lymph nodes of neck) | A | N/A | | | | 35.877 |

| | | | | | | | | |
|---|---|---|---|---|---|---|---|---|
| 13PM0931 | 73 | IV | Metastatic mucinous carcinoma (metastasis to lung) | A | N/A | | Yes | 30.148 |

*From date serum sample was taken to date of death

**Supplementary Table S3. Details for each of the breast cancer patients from the Circ.BR cohort used in this study.** Patient details were provided by the Brisbane Breast Bank with written informed consent from all patients.

| Specimen no. | Age | Breast cancer stage | Type of breast cancer | Vital status | Date of death | Survival time (years)* | Recurrence |
|---|---|---|---|---|---|---|---|
| 10-14-183 | 51 | T3N3a | Invasive ductal carcinoma with micropapillary carcinoma | D | 25/10/2016 | 2.2 | Yes |
| 10-13-139 | 74 | N/A | Inflammatory breast cancer | D | 22/04/2015 | 4.3 | Yes |
| 10-14-093 | 56 | T2N3a | Invasive ductal carcinoma | D | 05/08/2016 | 2.3 | Yes |
| 10-14-193 | 72 | T3N2a | Invasive ductal carcinoma | D | 04/01/2018 | 3.4 | Yes |
| 10-14-162 | 51 | T2N3a | Invasive ductal carcinoma | D | 17/11/2015 | 1.3 | Yes |
| 10-14-092 | 62 | T3Nx | Invasive ductal carcinoma | D | 07/12/2015 | 1.6 | Yes |
| 10-14-131 | 43 | T3N3a | Invasive lobular carcinoma | A | N/A | | Yes |
| 10-14-062 | 50 | T2N3a | Mixed invasive ductal carcinoma/micropapillary | A | N/A | | Yes |
| 10-16-028 | 69 | T2N1 | Mixed invasive ductal carcinoma/invasive lobular carcinoma | A | N/A | | Yes |
| 10-14-017 | 45 | T3N2 | Invasive ductal carcinoma | A | N/A | | No |
| 10-14-142 | 58 | T3N1a | Invasive ductal carcinoma | A | N/A | | No |
| 10-13-204 | 39 | T3N1MX | Invasive ductal carcinoma | A | N/A | | No |
| 10-13-186 | 53 | T1N1c | Invasive ductal carcinoma | A | N/A | | No |
| 10-13-157 | 36 | T3N1miM0 | Invasive ductal carcinoma | A | N/A | | No |
| 10-14-122 | 37 | T2N1a | Mixed metaplastic | A | N/A | | No |

*Time to death from date of diagnosis

**Figure S1**

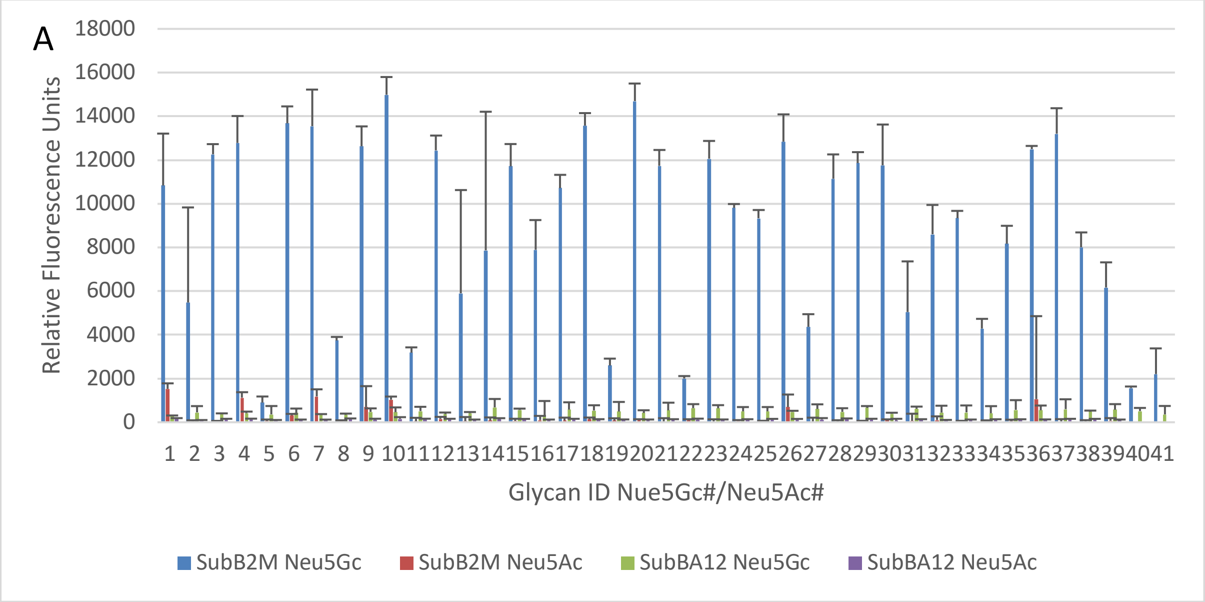
A
18000
16000
14000
12000
10000
8000
6000
4000
2000
0
Relative Fluorescence Units
1 2 3 4 5 6 7 8 9 10 11 12 13 14 15 16 17 18 19 20 21 22 23 24 25 26 27 28 29 30 31 32 33 34 35 36 37 38 39 40 41
Glycan ID Nue5Gc#/Neu5Ac#
SubB2M Neu5Gc
SubB2M Neu5Ac
SubBA12 Neu5Gc
SubBA12 Neu5Ac

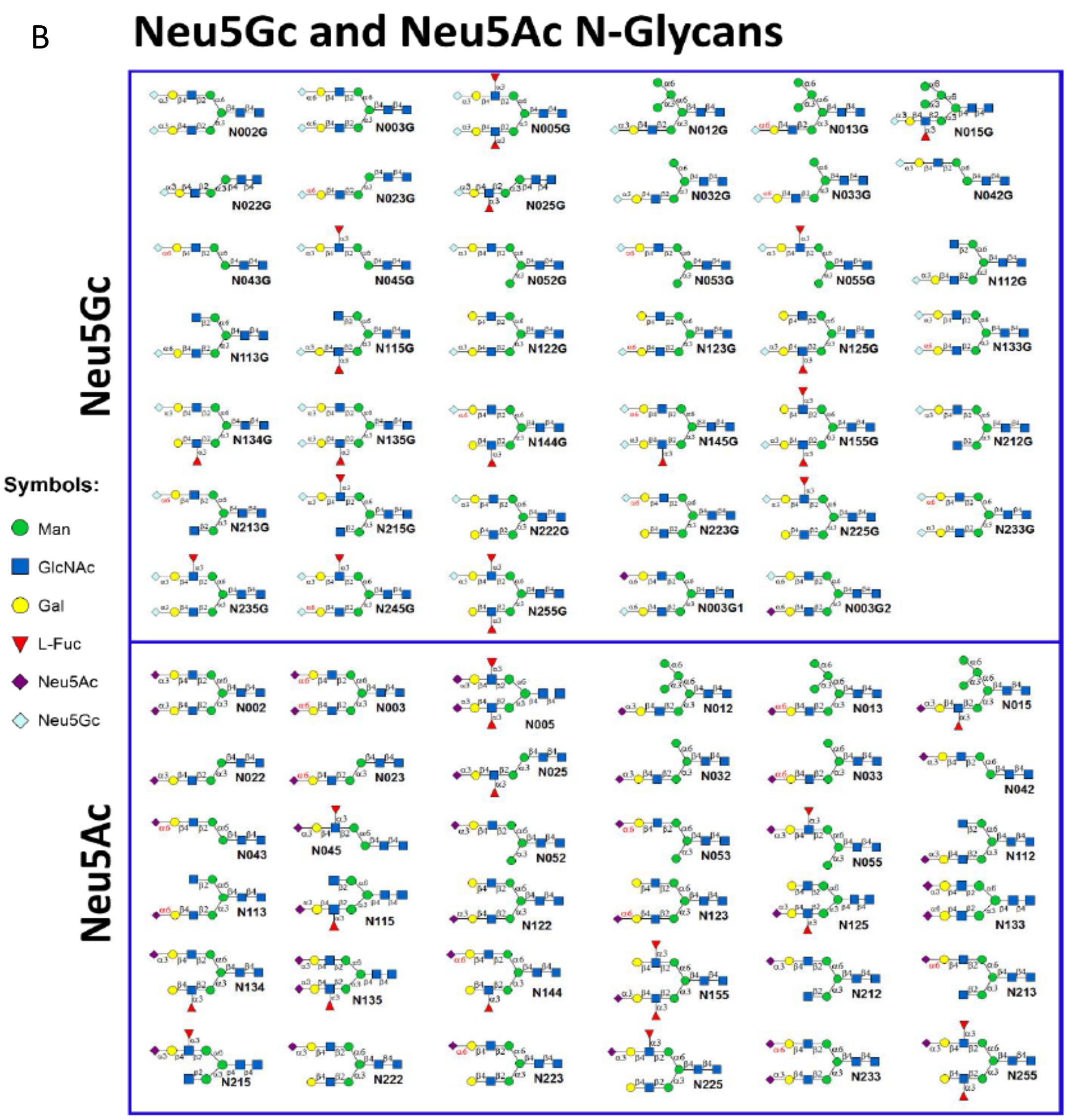
B
Neu5Gc and Neu5Ac N-Glycans
Neu5Gc
N002G
N003G
N005G
N012G
N013G
N015G
N022G
N023G
N025G
N032G
N033G
N042G
N043G
N045G
N052G
N053G
N055G
N112G
N113G
N115G
N122G
N123G
N125G
N133G
N134G
N135G
N144G
N145G
N155G
N212G
N213G
N215G
N222G
N223G
N225G
N233G
N235G
N245G
N255G
N003G1
N003G2
Neu5Ac
N002
N003
N005
N012
N013
N015
N022
N023
N025
N032
N033
N042
N043
N045
N052
N053
N055
N112
N113
N115
N122
N123
N125
N133
N134
N135
N144
N155
N212
N213
N215
N222
N223
N225
N233
N255
Symbols:
Man
GlcNAc
Gal
L-Fuc
Neu5Ac
Neu5Gc

### Figure S2

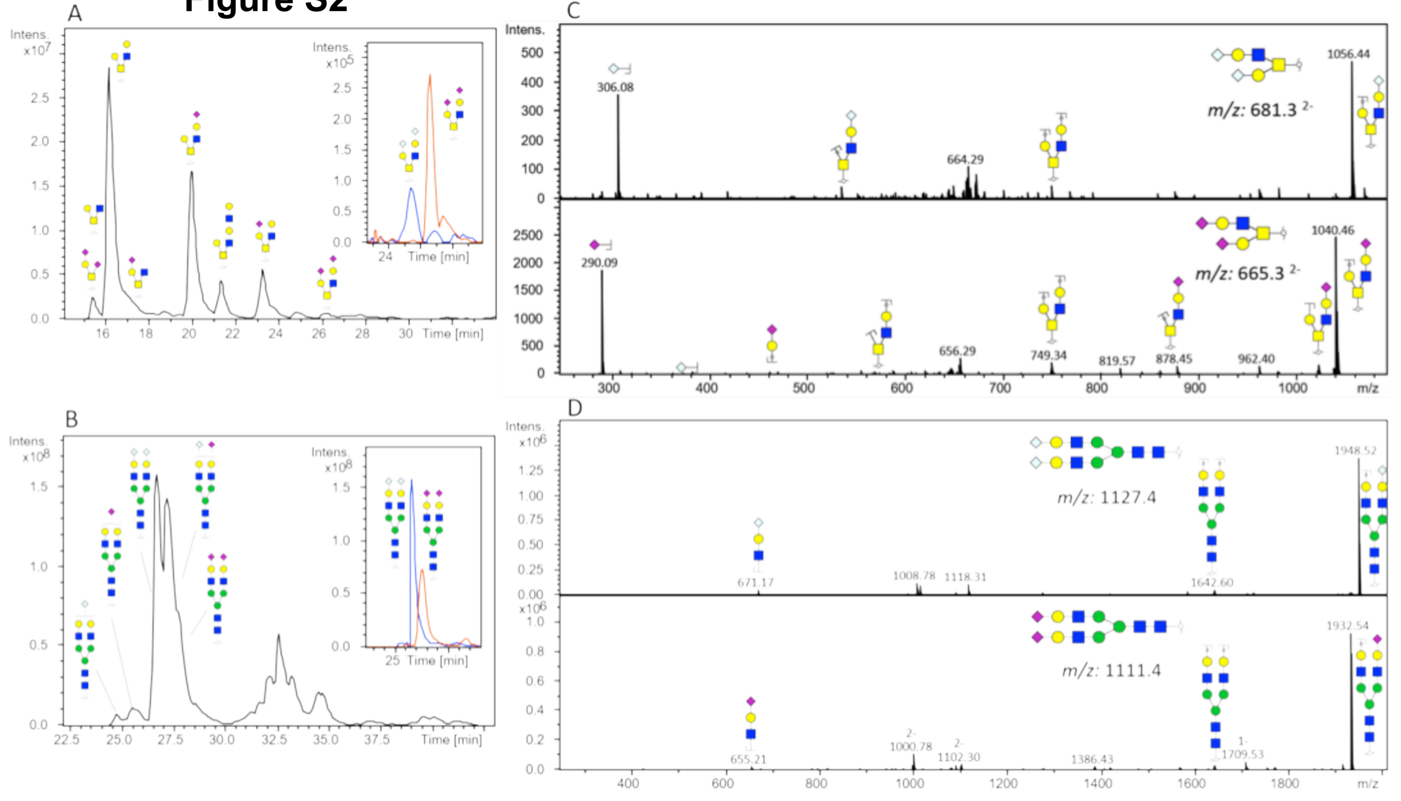
A
Intens. x10^7
Time [min]
B
Intens. x10^8
Time [min]
C
Intens.
306.08
664.29
1056.44
m/z: 681.3 2-
290.09
656.29
749.34
819.57
878.45
962.40
1040.46
m/z: 665.3 2-
m/z
D
Intens. x10^6
671.17
1008.78
1118.31
1642.60
1948.52
m/z: 1127.4
655.21
1000.78
1102.30
1386.43
1709.53
1932.54
m/z: 1111.4
m/z

**Figure S3**

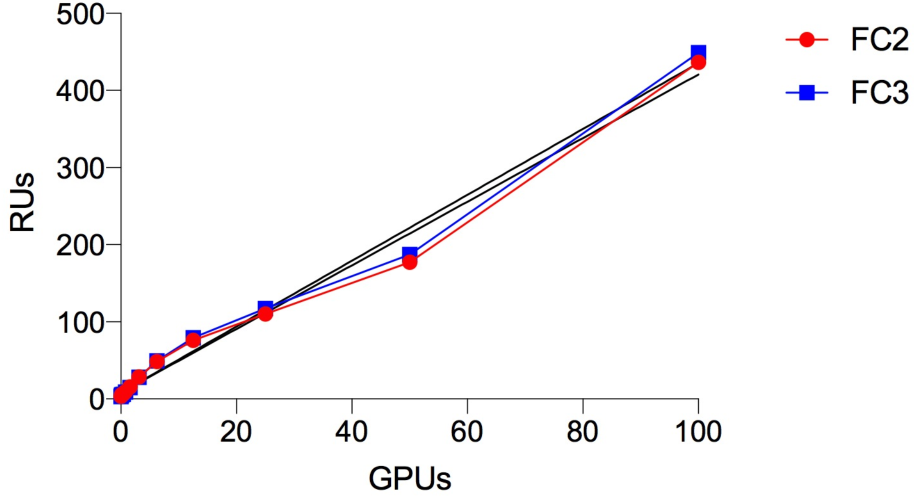

### Figure S4

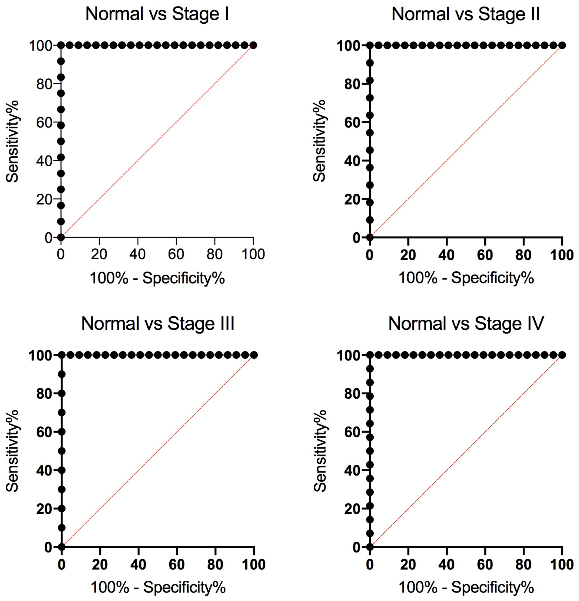

Normal vs Stage I
Normal vs Stage II
Normal vs Stage III
Normal vs Stage IV
Sensitivity%
100% - Specificity%
0
20
40
60
80
100

### Figure S5

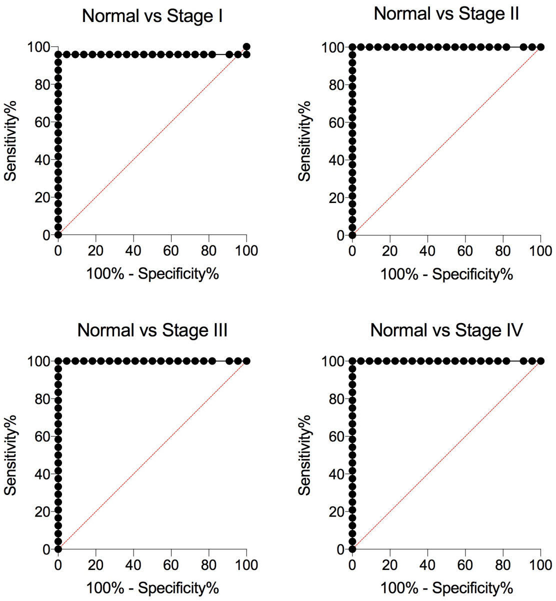

Normal vs Stage I
Normal vs Stage II
Normal vs Stage III
Normal vs Stage IV
Sensitivity%
100% - Specificity%
0
20
40
60
80
100

### Figure S6A

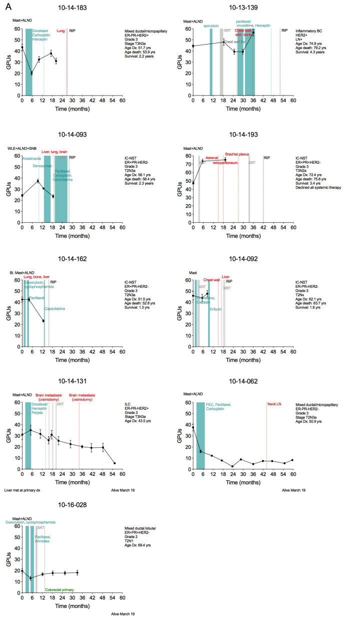

A
10-14-183
Mast+ALND
Docetaxel
Carboplatin
Herceptin
Lung
RIP
GPUs
Time (months)
Mixed ductal/micropapillary
ER-PR-HER2+
Grade 3
Stage T3N3a
Age Dx: 51.7 yrs
Age death: 53.9 yrs
Survival: 2.2 years
10-13-139
Mast+ALND
epirubicin
XRT
paclitaxel
vinorelbine, Herceptin
Chest wall,
skin, bone
Chest wall Bx
RIP
Inflammatory BC
HER2+
LN+
Age Dx: 74.9 yrs
Age death: 79.2 yrs
Survival: 4.3 years
10-14-093
WLE+ALND+SNB
Liver, lung, brain
Anastrozole
Denosumab
Paclitaxel,
Carboplatin,
Gemcitabine
XRT
RIP
IC-NST
ER+PR-HER2-
Grade 3
T2N3a
Age Dx: 56.1 yrs
Age death: 58.4 yrs
Survival: 2.3 years
10-14-193
Mast+ALND
Adrenal
retroperitoneum
Brachial plexus
XRT
RIP
IC-NST
ER+PR+HER2-
Grade 3
T3N2a
Age Dx: 72.4 yrs
Age death: 75.8 yrs
Survival: 3.4 yrs
Declined all systemic therapy
10-14-162
Bi. Mast+ALND
Lung, bone, liver
Doxorubicin,
cyclophosphamide
Paclitaxel
Capecitabine
RIP
IC-NST
ER+PR-HER2-
Grade 3
T2N3a
Age Dx: 51.5 yrs
Age death: 52.8 yrs
Survival: 1.3 yrs
10-14-092
Mast
Chest wall
Liver
RIP
XRT
Eribulin
IC-NST
ER-PR-HER2-
Grade 3
T2Nx
Age Dx: 62.1 yrs
Age death: 63.7 yrs
Survival: 1.6 yrs
10-14-131
Mast+ALND
Brain metastasis
(crainiotomy)
Docetaxel
Herceptin
Perjeta
XRT
ILC
ER-PR-HER2+
Grade 3
Stage T3N3a
Age Dx: 43.5 yrs
Liver met at primary dx
Alive March 19
10-14-062
Mast+ALND
FEC, Paclitaxel,
Carboplatin
Neck LN
Mixed ductal/micropapillary
ER-PR-HER2-
Grade 3
Stage T2N3a
Age Dx: 50.9 yrs
Alive March 19
10-16-028
Mast+ALND
Doxorubicin, cyclophosphamide
XRT
Paclitaxel,
Arimidex
Colorectal primary
Mixed ductal lobular
ER+PR+HER2-
Grade 2
T2N1
Age Dx: 69.4 yrs
Alive March 19

### Figure S6B

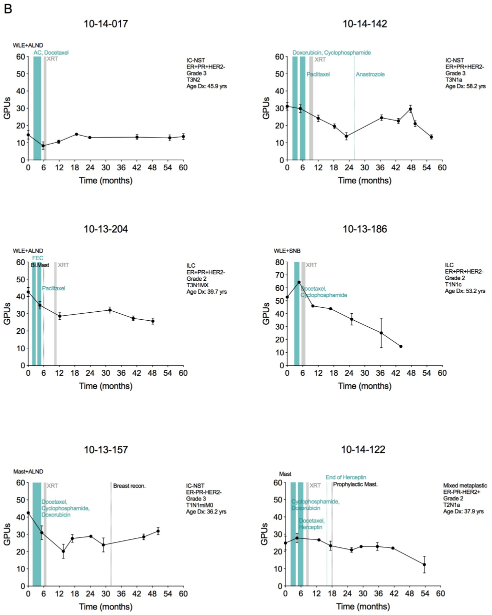

B
10-14-017
WLE+ALND
AC, Docetaxel
XRT
IC-NST
ER+PR+HER2-
Grade 3
T3N2
Age Dx: 45.9 yrs
GPUs
Time (months)
10-14-142
Doxorubicin, Cyclophosphamide
XRT
Paclitaxel
Anastrozole
IC-NST
ER+PR+HER2-
Grade 3
T3N1a
Age Dx: 58.2 yrs
GPUs
Time (months)
10-13-204
WLE+ALND
FEC
Bl.Mast
XRT
Paclitaxel
ILC
ER+PR+HER2-
Grade 2
T3N1MX
Age Dx: 39.7 yrs
GPUs
Time (months)
10-13-186
WLE+SNB
XRT
Docetaxel,
Cyclophosphamide
ILC
ER+PR+HER2-
Grade 2
T1N1c
Age Dx: 53.2 yrs
GPUs
Time (months)
10-13-157
Mast+ALND
XRT
Breast recon.
Docetaxel,
Cyclophosphamide,
Doxorubicin
IC-NST
ER-PR-HER2-
Grade 3
T1N1miM0
Age Dx: 36.2 yrs
GPUs
Time (months)
10-14-122
Mast
End of Herceptin
XRT
Prophylactic Mast.
Cyclophosphamide,
Doxorubicin
Docetaxel,
Herceptin
Mixed metaplastic
ER-PR-HER2+
Grade 2
T2N1a
Age Dx: 37.9 yrs
GPUs
Time (months)
